# Effects of Chitosan as a Permeabilizing Agent in Different Yeast Species. Studying Enzymes *in situ*

**DOI:** 10.64898/2026.05.06.723273

**Authors:** Minerva Araiza-Villanueva, Norma Silvia Sánchez, Martha Calahorra, Francisco Padilla-Garfias, Antonio Peña

**Affiliations:** Departamento de Genética Molecular, Instituto de Fisiología Celular, Universidad Nacional Autónoma de México, Circuito Exterior s/n, Ciudad Universitaria, 04510 México City, México

**Keywords:** Chitosan, Yeast, Plasma membrane permeabilization, *In situ* enzyme activity

## Abstract

Chitosan is an oligosaccharide derived from chitin that is protonated at acidic pH to form a polycation. Its positive charge promotes the interaction with negatively charged components of the yeast cell surface, which has been associated with increased cell permeability and growth inhibition. In this study, we investigated the interaction of chitosan with the cell surface and its permeabilizing capacity in three yeast species displaying distinct susceptibility profiles, *Saccharomyces cerevisiae*, *Candida albicans* and *Debaryomyces hansenii*. We evaluated the correlation between differential susceptibility and chitosan association at the cell surface, as well as cell permeabilization, by integrating growth analyses with surface-binding assays, including FITC-conjugated chitosan to monitor surface association and cellular integration over time, and ultrastructural examination by transmission electron microscopy (TEM). Our results showed that chitosan exhibited varying effects on the growth and permeability of each yeast strain, with *D. hansenii* being the most susceptible. Furthermore, we observed the incorporation of chitosan onto the cell surface and confirmed its role as a permeabilizing agent. Finally, we used chitosan-induced permeabilization as a method to measure the activity of selected enzymes *in situ*, demonstrating its potential for studying metabolic functions in permeabilized yeast cells. Overall, our findings establish chitosan as a strain-dependent antifungal agent and a useful tool for functional biochemical analyses in yeast.

## 1. Introduction

Chitosan is a linear polycationic polysaccharide derived from the deacetylation of chitin, mainly composed of glucosamine (GlcN) and to a lesser extent by N-acetylglucosamine (GlcNAc) (Aranaz et al., 2009; Lopez-Moya et al., 2019). In aqueous solution, its solubility increases under acidic conditions, due to the protonation of free amino groups (Aranaz et al., 2021; Pal et al., 2021).

Chitosan has been widely used in the production of nanomaterials, as a flocculant, antioxidant and antimicrobial agent, and is effective against a broad range of bacteria, filamentous fungi and yeasts (Lopez-Moya et al., 2019; Pal et al., 2021). Importantly, it is harmless to mammalian cells at concentrations that are fungicidal to human pathogenic fungi, highlighting its potential as a natural antifungal agent suitable for clinical applications (Lopez-Moya et al., 2019).

In filamentous fungi, chitosan has demonstrated a variety of antifungal mechanisms. In *Rhizopus stolonifer,* concentrations of 2.0 mg/mL caused plasma membrane alterations and potassium (K^+^) leakage, as well as the inhibition of the plasma membrane H^+^-ATPase, indicating that the effect of chitosan on *R. stolonifer* is likely due to destabilization of the plasma membrane structure (García-Rincón et al., 2010). In *Aspergillus ochraceous*, chitosan inhibited spore germination and mycelial growth in a dose-dependent manner, also inducing changes in the morphology of the hyphae, and plasma membrane fluidity by altering phospholipid and sterol metabolism. It has also been suggested that chitosan may impact the cell wall, whose integrity could be disrupted through the expression of cell wall hydrolases (Meng et al., 2020).

Aranda-Martinez et al., (2016) found that in the yeast *Candida albicans*, chitosan exhibits an antifungal effect by disrupting the plasma membrane and altering cellular homeostasis. At low concentrations (2.5 – 10 μg/mL), it induces potassium efflux, acidification of the external medium and an increased calcium uptake; however, large molecules such as proteins remain inside, and retain their functionality. It was further observed that respiration was partially restored after the addition of phosphate, MgCl_2_, ATP and NAD^+^, indicating that, although the cell surface is damaged, the remaining internal contents of the cell remain functional. At moderate concentrations (20 – 50 μg/mL) it inhibits fermentation and respiration, and at higher concentrations (> 1 mg/mL), it causes membrane damage, leading to ionic imbalance, metabolic disruption, and growth inhibition (Peña et al., 2013). Besides, low concentrations of chitosan also reduce biofilm formation in *C. albicans*, without killing all the cells (Costa et al., 2014).

Another study in *C. albicans* showed the effects of high molecular weight chitosan oligosaccharides (HCOS), and reported that they inhibit cell growth, and disrupt cell wall integrity by interfering with the activity of Phr1p, a critical enzyme involved in β-glucan elongation and branching. By contrast, low molecular weight chitosan oligosaccharides (LCOS) exhibited significantly reduced antifungal activity compared to HCOS. This difference is attributed to HCOS’s ability to integrate into the fungal cell wall, whereas LCOS are primarily localized inside the cell. Furthermore, HCOS display synergistic activity with fluconazole, effectively inhibiting biofilm formation (Li et al., 2022).

In the human pathogen, *Histoplasma capsulatum*, chitosan also exhibits a significant antifungal activity in both the filamentous and the yeast form, the latter being the most susceptible. In this study, chitosan was used along with some antifungal drugs such as itraconazole, enhancing its antifungal effect and altering biofilm formation, and it was also found to induce oxidative stress (Brilhante et al., 2023).

The antifungal effects of chitosan vary among fungal species, depending on differences in plasma membrane and cell wall composition, developmental stage, and environmental conditions.(Poznanski et al., 2023). While the plasma membrane and cell wall are recognized as the primary targets of chitosan, its impact extends beyond these structures. For instance, in *Neurospora crassa*, transcriptomic analysis of cells exposed to chitosan at different time points revealed an overexpression of genes related to oxidative stress, including glutathione transferases (GST-1, GST-4) and dioxygenases, suggesting that this polymer elicits oxidative stress as a part of its multifaceted mechanism of action (Lopez-Moya et al., 2016). These multiple mechanisms reinforce the potential of chitosan as a broad-spectrum antifungal agent. Moreover, a more integrated understanding of these mechanisms is necessary to fully elucidate chitosan’s mode of action across various fungi.

Given these findings, the effects of chitosan were investigated in three yeast species that displayed distinct susceptibility profiles to chitosan, including *Saccharomyces cerevisiae*, *Candida albicans*, and *Debaryomyces hansenii*.

Specifically, we explored the interaction of chitosan with the yeast cell surface and evaluated its cell-permeabilizing capacity. This comparative approach allowed us to explore whether differences in susceptibility are associated with variations in surface binding and permeabilization. Furthermore, we used the membrane-permeabilizing properties of chitosan to facilitate the *in-situ* analysis of selected enzyme activities, providing functional and biochemical insights into the effects of chitosan across strains.

## 2. Materials and Methods

### 2.1 Strains and growth conditions

Experiments were carried out on the *C. albicans* strain ATCC 10231, *S. cerevisiae* strain ATCC S288C, and *D. hansenii* strain Y7426 (also known as NRRL Y-7426 or CBS 767). Cells were maintained in solid YPD medium (1% yeast extract, 2% peptone, 2% glucose and 2% agar), renewing the cultures once a month.

Cells were grown in YPD or YNBG (Yeast nitrogen base w/o amino acids from Bio Basic Inc., 2% glucose) liquid media, with or without chitosan, in an orbital shaker at 250 rpm and 30 °C.

To determine the susceptibility to chitosan (section 2.4), yeast cells were grown in RPMI-1640 medium supplemented with L-Glutamine and phenol red, without bicarbonate (Thermo Fisher) and buffered with morpholine propane sulfonate (MOPS) to pH 7.0 (Rex, 2008).

For fasting conditions, cells grown for 24 h in YPD medium were collected by centrifugation at 5524 x g and washed twice with distilled water, then resuspended in 250 mL of distilled water and starved for 24 h for *S. cerevisiae* S288C, or 48 h for *C. albicans* and *D. hansenii*.

### 2.2 Chitosan preparation and FITC–chitosan preparation

Low molecular weight chitosan (MW = 50 - 190 kDa, degree of deacetylation 75 – 85%, Sigma Aldrich) was dissolved at a final concentration of 10 mg/mL in 0.1 M lactic acid, to reach a final pH of 4.5.

Fluorescein isothiocyanate (FITC; 10 mg) was dissolved in 10 mL of anhydrous ethanol and added dropwise to 10 mL of the chitosan solution (10 mg/ mL) under constant stirring. The reaction mixture was incubated in the dark at 4 °C overnight (∼16-18 h) to allow covalent coupling between FITC isothiocyanate groups and primary amines of chitosan. After incubation, pH was adjusted to 10–11 by addition of 1 M NaOH, inducing the formation of a gel-like precipitate. The suspension was centrifuged at 2000 x g, and the resulting pellet was washed twice with anhydrous ethanol to remove unreacted FITC, followed by two washes with Milli-Q water. The material was then centrifuged and filtered through sterile gauze to eliminate excess liquid. The resulting FITC-chitosan (FITC-CTS) gel was protected from light and dried by lyophilization for 48 h. Finally, a 10 mg/mL stock solution of FITC-CTS was prepared with acetic acid 0.1 M (Li et al., 2022; Moussa et al., 2013).

### 2.3 Growth assays under chitosan treatment

Yeast growth was followed to evaluate the effects of increasing chitosan concentrations on yeast proliferation under liquid and solid growth conditions. For experiments in liquid media, precultures grown for 18 h in YPD medium were harvested and washed twice with distilled water, then adjusted to an optical density (OD_600nm_) of 0.03 units in liquid YNBG medium containing chitosan concentrations ranging from 0 to 5000 μg/mL for *S. cerevisiae* and *C. albicans*, and from 0 to 5 μg/mL for *D. hansenii* (as it showed to be more sensitive to higher concentrations of chitosan). Growth was monitored using a BioScreen C automated plate reader in 100-well plates at 30 °C (or 28°C for *D. hansenii*) for 72 h, recording absorbance at 600 nm every 30 min, with constant shaking.

For growth assessment on solid YNBG medium, precultures grown for 18 h were harvested, washed twice with distilled water and resuspended to an OD_600nm_ of 1.0. Cell suspensions were distributed in a 96-well plate, and six consecutive 1:10 serial dilutions were prepared. Aliquots from each dilution were spotted onto YNBG agar plates containing chitosan at concentrations ranging from 0 to 1000 μg/mL using a pin replicator, following a drop test assay format. This concentration range was selected because higher chitosan concentrations compromised agar consistency and hindered proper plate solidification. Plates were incubated on 30 °C for 72 h, and colony growth was documented at day 3.

### 2.4 Determination of MIC and MFC

Yeast tolerance to chitosan was assessed using a broth microdilution method, according to the Clinical & Laboratory Standards Institute (CLSI) guidelines (M27-A3) (Rex, 2008), with modifications. For this, 24 h precultures grown in YPD medium were washed twice with distilled water and adjusted to a McFarland standard of 0.5 (OD_600nm_ 0.1 U/mL) in RPMI medium containing final chitosan concentrations ranging from 0 to 5000 μg/mL. Growth was monitored using a BioScreen C automated plate reader in 100-well plates at 37 °C. For *D. hansenii,* which is sensitive to temperatures above 30 °C, assays were performed at its optimal temperature of 28 °C.

Growth inhibition was quantified by calculating the area under the curve (AUC) from optical density (OD) measurements at 600 nm over time instead of visual inspection. The percentage of growth inhibition was calculated using the following equation:

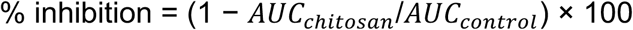

The inhibition values were plotted as a function of chitosan concentration and fitted using nonlinear regression in GraphPad Prism. The minimum inhibitory concentration (MIC) was defined as the lowest chitosan concentration resulting in ≥ 50% reduction of the AUC compared to the untreated control. The minimum fungicidal concentration (MFC) was determined as described by Pfaller and Fontenelle (Fontenelle et al., 2007; Pfaller, 2005). Briefly, aliquots (10 μL) from the concentrations where no visible growth appeared after 48 h were plated on potato dextrose agar (PDA) and incubated for an additional 48 h at 30 °C. The MFC was defined as the lowest concentration yielding ten or fewer colonies (Fontenelle et al., 2007; Li et al., 2022; Pfaller, 2005; Rex, 2008).

### 2.5 Cell viability

Colony forming units (CFU) were determined to evaluate cell viability after treatment with different concentrations of chitosan. Cells previously grown in YPD liquid medium for 24 h were washed twice with distilled water and incubated for 10 min at 30 °C in YNBG medium with different concentrations of chitosan at 1 × 10^6^ cells/mL in a final volume of 1 mL. Then, 100 μL of a 1,000-fold dilution (approximately 100 cells) was spread on solid YPD medium, and colonies were counted after 48 h.

### 2.6 Transmission electron microscopy

The effects of chitosan on yeast cells were analyzed by transmission electron microscopy (TEM). Yeast cells were cultured in YPD medium for 24 h, harvested by centrifugation, washed twice with sterile water and resuspended to 50% (wet weight/ volume). Cell suspensions were normalized to a final concentration of 10 mg/mL and treated with strain-specific chitosan concentrations selected based on the known tolerance of each yeast species, to evaluate ultrastructural alterations across a low-to-high concentration range. Accordingly, *S. cerevisiae* cells were treated with 10 and 500 μg/mL chitosan, *C. albicans* with 5 and 500 μg/mL, and *D. hansenii* with 0.5 and 20 μg/mL for 10 min. After treatment, the cells were washed again, collected by centrifugation and the supernatant was discarded. The resulting pellets were fixed with 2.5% glutaraldehyde using a modified protocol based on the previously described by Padilla-Garfias et al. (2022). Images were captured using a JEOL-JEM-1200 EXII transmission electron microscope.

### 2.7 Zeta potential determination

Starved cells were placed in a medium containing 10 mM MES-TEA buffer (2-(N-morpholino) ethanesulfonic acid adjusted to pH 6.0 with triethanolamine), 2 mM glucose, 1 mg/mL of starved cells (0.1% wet weight), and chitosan at concentrations of 0, 0.25, 0.5, 2.5, 10, and 100 μg/mL (final volume of 13 mL).

The displacement velocity of these cells under an electric field was then measured directly under the microscope using a Zeta-Meter ZM-75. This measurement was used to calculate the zeta potential according to the Helmholtz–Smoluchowski equation:

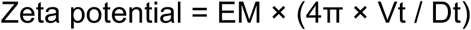

where EM is the electrophoretic mobility; Vt is the viscosity of the liquid, measured in poises at the temperature of measurement; Dt is the dielectric constant of the suspending fluid at the temperature of measurement; and 4π = 12.57.

It should be noted that, in comparison with the experiments used to determine cell permeabilization such as propidium iodide (PI) uptake and potassium efflux, the amounts of wet cell weight and chitosan concentration required were 10-fold higher; however, the cell-to-chitosan ratio was maintained.

### 2.8 FITC-chitosan Binding to the Cell Surface

To assess chitosan binding to the cellular surface, yeast cells were prepared at a concentration of 10 mg/mL in a solution containing 10 mM MES-TEA (pH 6.0) and a final concentration of 10 μg/mL of FITC-chitosan (FITC–CTS).

Samples were incubated in the dark for 10 min, 1 h, 2 h, 4 h, and 24 h. At each time point, 5 μL aliquots were placed onto glass slides for fluorescence microscopy analysis. Control conditions included untreated cells (negative control) and cells treated with FITC alone (5 μg/mL).

Cells were visualized at 100x using an Olympus BX51 microscope equipped with Ocular Image Acquisition Software (QImaging). Images were subsequently analyzed using ImageJ software.

### 2.9 Determination of membrane integrity

Membrane integrity assay was performed using PI, in different yeast strains treated with chitosan. Briefly, yeast cells (50 mg, wet weight) were starved as required and incubated in test tubes in a final volume of 5.0 mL (10 mg/mL) containing 10 mM MES-TEA buffer (pH 6.0), 20 mM glucose, 5 μg/mL PI, and varying concentrations of chitosan. Incubation was carried out at 30 °C for 20 minutes. A control for 100% membrane permeabilization was prepared by boiling a similar cell suspension in a water bath for 20 minutes, followed by cooling on ice and staining with 5 μg/mL PI. After incubation or boiling, fluorescence intensity was measured (excitation wavelength: 535 nm; emission wavelength: 620 nm) using a FLUOstar Omega plate reader. Cells were then observed under an Olympus BX51 microscope, images were acquired at 40x magnification using Ocular Image Acquisition Software (QImaging) and further processed with ImageJ software.

### 2.10 Measurement of Potassium efflux

To measure potassium efflux, a K^+^ selective electrode was used. For this, 100 mg wet weight of fasted cells were incubated in 2 mM MES-TEA buffer (pH 6.0) and 20 mM glucose in a final volume of 10 mL. The potassium changes in the medium were monitored by varying the concentrations of chitosan added to the cells. The experiments were carried out in a chamber under continuous magnetic stirring at 30°C.

### 2.11 NADH uptake as an indicator of membrane permeability

NADH uptake was indirectly assessed by measuring the reduction of the vital dye Alamar Blue (Invitrogen). This assay was designed based on the principle that if chitosan treatment affects membrane permeability, charged molecules that lack plasma membrane transporters, such as NADH, can enter the cell. The NADH-mediated reduction of Alamar Blue’s resazurin to resorufin generates fluorescence, which serves as an indicator of membrane permeability. In addition, permeabilized cells retain enzymes, such as dehydrogenases, capable of catalyzing redox reactions involving NADH and resazurin.

For this assay, yeast cells were grown in YPD medium for 24 h and subsequently starved for 24 or 48 h, as indicated. Cell suspensions were normalized to 10 mg/mL and treated with different concentrations of chitosan for 10 min, washed twice, and then adjusted to an OD_600nm_ of 0.5 in a solution containing 10 mM MES-TEA buffer (pH 6.0), 20 mM glucose, and 0.8 M sorbitol to prevent osmotic stress. NADH was added to a final concentration of 60 μg/mL. The reaction was carried out in 96-well plates and incubated at 30 °C for 30 min. Fluorescence was measured using a FLUOstar Omega plate reader (excitation wavelengths: 530-560 nm, emission wavelength: 590 nm) according to the manufacturer’s instructions. Boiled cells were used as negative controls and untreated cells as non-permeabilized controls. Fluorescence data were normalized to the untreated control.

### 2.12 Preparation of permeabilized cells and cytoplasmic extracts for enzymatic measurements

Cells were cultured in YPD medium and starved as previously described, then collected by centrifugation and resuspended in distilled water to a ratio of 0.5 g/mL (wet weight). These cells were permeabilized as previously indicated in 10 mM MES-TEA buffer pH 6.0, with varying concentrations of chitosan (depending on the strain) in a final volume of 10 mL. The suspension was incubated in a tube rotator at 30 °C for 15 minutes. After incubation, the cells were centrifuged to remove chitosan, and the cells were carefully washed with distilled water to eliminate residual chitosan. The permeabilized cells were resuspended in a solution of 0.8 M sorbitol to prevent osmotic stress, and subsequently used to determine the activity of various enzymes, as described in section 2.13.

For cytoplasmic extracts, the same permeabilized cells were used. Cells were washed twice with PBS buffer (pH 7.0) and resuspended in lysis buffer (100 mM HEPES/KOH, pH 7.5; 600 mM potassium acetate; 10 mM magnesium acetate; 1 mM EDTA; 20% glycerol; 8 mM 2-mercaptoethanol; and a protease inhibitor tablet (’Complete’, Roche). An equal volume of 0.5 mm diameter glass beads was added to the cell suspension. The cells were mechanically disrupted using cycles of vortexing (60 seconds) and ice cooling (120 seconds) for a total of 10 cycles. The homogenate was then collected without centrifugation, separating the precipitated beads from the supernatant. The homogenate was used to measure enzymatic activity.

### 2.13 Measurement of enzyme activities

The enzymatic activities of alcohol dehydrogenase (ADH), glucose-6-phosphate dehydrogenase (G6PD), hexokinase (HK), glutamate dehydrogenase (GDH) and pyruvate kinase (PK) (Hess and Wieker, 1974; Sánchez et al., 2008, 2006) were determined in both permeabilized cells and cytoplasmic extracts. Absorbance changes were measured using an OLIS CLARITY UV/Vis spectrophotometer. This instrument allowed accurate measurement of absorbance in both light-scattering (turbid or hazy) samples and clear solutions, ensuring precise detection of enzymatic activity. Enzymatic activity was normalised to milligrams of protein (Markwell et al., 1978).

### 2.14 Statistical Analysis

Statistical analyses were carried out using GraphPad Prism version 10.0.0 (GraphPad Software, Boston, MA, USA; www.graphpad.com).

For comparisons involving multiple treatments against a single control, one-way or two-way analysis of variance (ANOVA) followed by Dunnett’s multiple comparisons test was applied, as required. For comparisons among multiple groups, two-way ANOVA followed by Tukey’s multiple comparisons test was used.

In experiments evaluating the effect of chitosan concentration within each yeast strain, analyses were performed by comparing treatments to the corresponding untreated control. For experiments involving two independent variables (strain and chitosan concentration), two-way ANOVA was used to assess the main effects. Differences were considered statistically significant at α = 0.05. Statistical significance is indicated as follows: *p < 0.05, **p < 0.01, ***p < 0.001, and ****p < 0.0001; ns, not significant.

## 3. Results

### 3.1 Effect of chitosan on yeast growth in liquid and solid media

Growth was initially evaluated in both YPD and YNBG liquid media, observing that cells grown in YPD displayed a markedly higher tolerance to chitosan, consistent with previous reports (Peña et al., 2013) (Supplementary figure 1). Based on these observations, subsequent experiments were performed using the minimal medium YNBG, in which the effect of chitosan on growth was more evident.

In *S. cerevisiae*, a concentration-dependent inhibitory effect on growth in liquid medium was observed starting at 5 μg/mL, while at 1000 μg/mL, there was a reduction of more than 50% in the area under the growth curve (AUC), compared with the untreated control (Figure 1a, Supplementary table 1). By contrast, growth on solid medium was largely preserved at 1000 μg/mL of chitosan, with no marked differences between this concentration and the untreated condition in solid medium (Figure 1b). This discrepancy may reflect differences in chitosan accessibility or diffusion between liquid and solid media.

**Figure 1.**
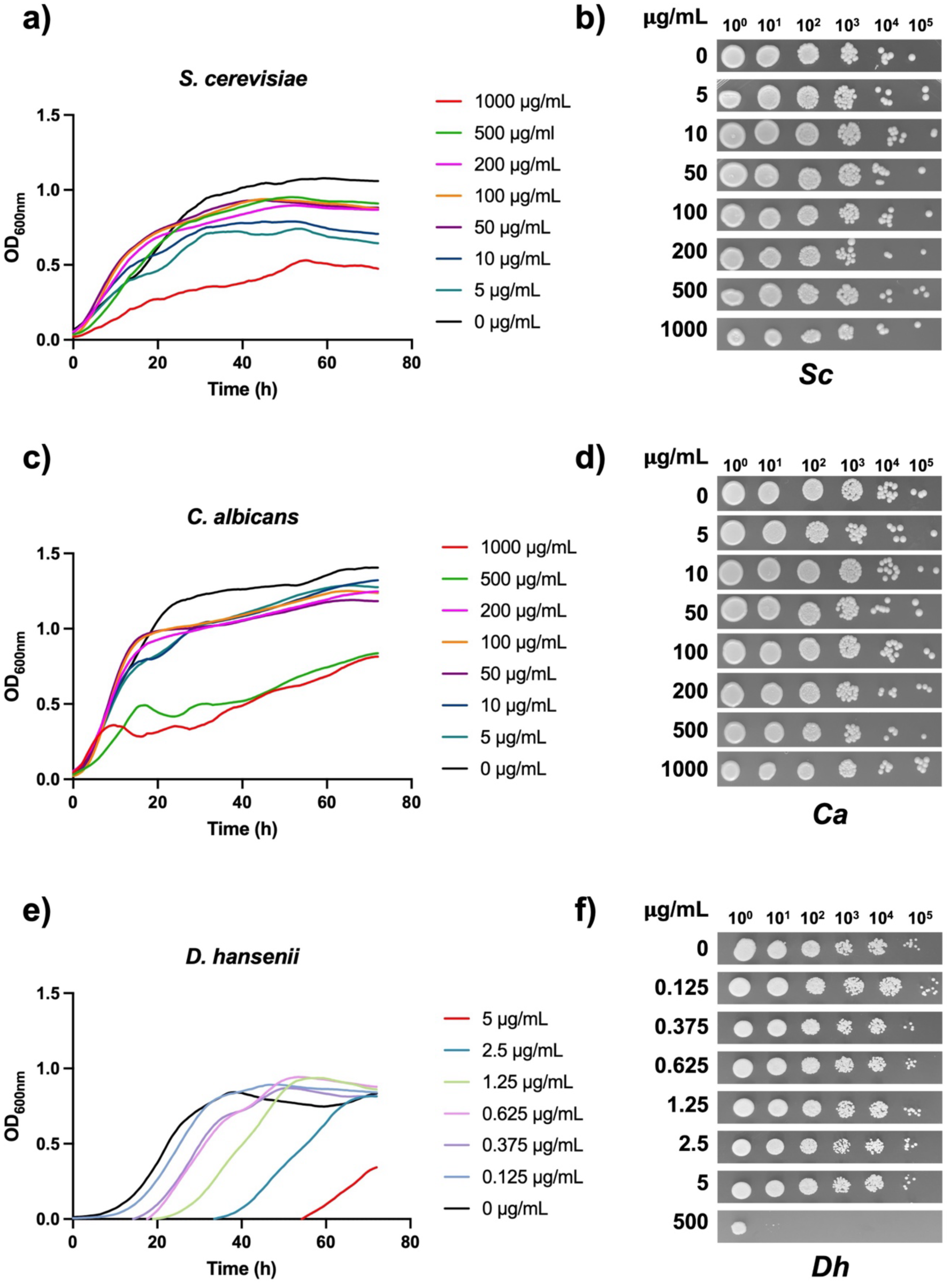
Effect of chitosan on the growth of yeast species. Growth curves of *S. cerevisiae* (a), *C. albicans* (c) and *D. hansenii* (e) in YNBG medium supplemented with different concentrations of chitosan. Curves represent the mean of three biological replicates, each with two technical replicates. Spot assays of *S. cerevisiae* (*Sc*) (b), *C. albicans* (*Ca*) (d) and *D. hansenii* (*Dh*) (f) were performed on solid YNBG medium containing different concentrations of chitosan and incubated for 72 h at 30 °C. Serial tenfold dilutions (six dilution steps) were prepared from cell suspensions adjusted to an (OD_600nm_)/mL of 1 unit, and spots were plated accordingly. Spot assays were performed using three biological replicates.

Also, the growth of *C. albicans* on a liquid YNBG medium supplemented with chitosan was more affected, with a reduction from 500 μg/mL, the area under the AUC decreases by more than 50% (Figure 1c, Supplementary table 1). As with *S. cerevisiae*, the growth of *C. albicans* on solid YNBG medium was unaffected by a concentration of 1000 μg/mL (Figure 1d).

Finally, *D. hansenii* exhibited the highest sensitivity to chitosan. Growth was markedly delayed in liquid YNBG medium, although the exponential growth rate appeared largely unaffected. The primary effect was observed during the lag phase, which was progressively extended as chitosan concentration increased, with the most pronounced delay at 5 µg/mL (Figure 1e). This behaviour is consistent with the AUC analysis, which revealed a clear reduction in total growth at concentrations of 5 µg/mL or higher. Specifically, the AUC decreased from ∼41.2 to 4.3 at 5 μg/mL. At higher concentrations (10-100 μg/mL), AUC values remained minimal (2.0-3.4), confirming a strong inhibitory effect on overall biomass accumulation despite eventual exponential growth (Supplementary table 1).

As observed for the other yeast species, tolerance to chitosan was higher in solid medium, with detectable growth even at 5 µg/mL. However, unlike *S. cerevisiae* and *C. albicans*, *D. hansenii* showed visible growth impairment at higher concentrations, particularly at 500 µg/mL (Figure 1f).

The growth patterns discerned among the three yeast species investigated indicate that chitosan manifests a fungistatic rather than a fungicidal one. This assertion is supported by the delayed growth recorded in the three strains, even at elevated concentrations of chitosan, rather than a decline in biomass indicative of cellular death.

### 3.2 Minimum inhibitory and fungicidal concentrations of chitosan differ among yeast species

As different levels of susceptibility to chitosan were observed among the yeast species, it was necessary to determine the minimum inhibitory concentration (MIC) and the minimum fungicidal concentration (MFC) of chitosan for each species. To determine chitosan tolerance across the different yeast species, MIC and MFC values were calculated based on growth inhibition and viability assays. The overall experimental workflow used for these determinations is summarized in Figure 2a. MIC values were calculated from nonlinear regression analysis of the percentage of growth inhibition data (Supplementary table 2). The MFC of chitosan for *S. cerevisiae* was estimated to be >5000 µg/mL, whereas MIC was 297 μg/mL. The exact MFC could not be determined, because concentrations above this value exceeded the maximum permitted for liquid culture in order to maintain proper buffering conditions and avoid growth inhibition caused by pH changes rather than by chitosan itself. In the case of *C. albicans*, the MIC and MFC values were 379 μg/mL and >5000 μg/mL, respectively. In contrast, *D. hansenii* exhibited markedly lower MIC and MFC values, with 0.8 μg/mL for MIC and 10 μg/mL for MFC. The MFC/MIC was higher than 4 in all species, indicating that chitosan exerted a fungistatic rather than fungicidal effect under the tested conditions (Figure 2b) (Dudiuk et al., 2019).

**Figure 2.**
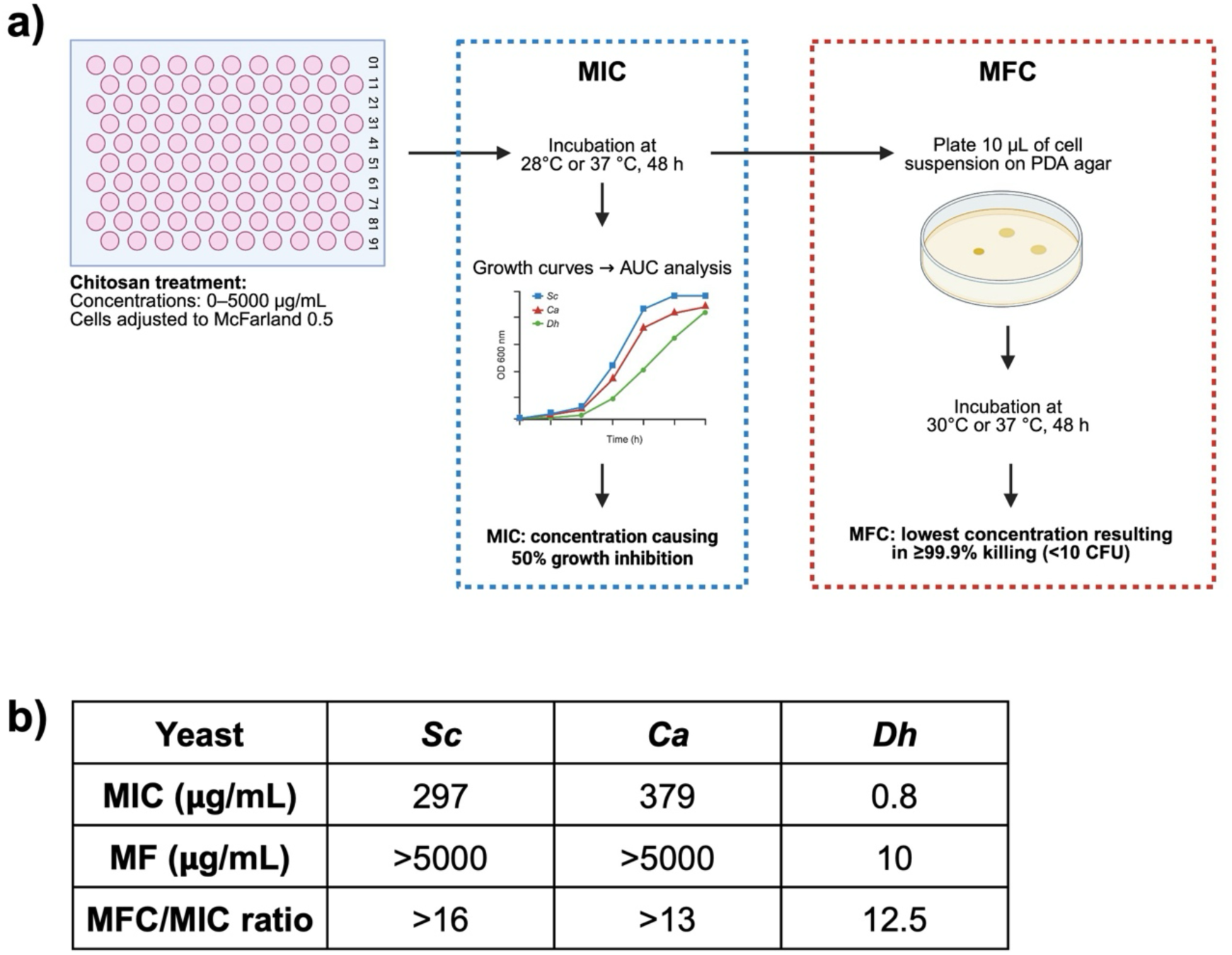
Effect of chitosan on cell viability of different yeast species. (a) Schematic workflow for MIC and MFC determination. Yeast cells were exposed to 0–5000 μg/mL chitosan. MIC was defined as the concentration causing 50% growth inhibition based on AUC analysis, whereas MFC was defined as the lowest concentration resulting in 99.9% killing (1-log CFU reduction). (b) Determination of chitosan tolerance by estimating the minimum inhibitory concentration (MIC) and minimum fungicidal concentration (MFC) in *S. cerevisiae* (*Sc*), *C. albicans* (*Ca*), and *D. hansenii* (*Dh*). MIC values were calculated from growth curves using area under the curve (AUC) and percentage inhibition analysis, as detailed in Supplementary Table 2. MFC was determined by plating treated cells and assessing colony-forming units (CFU). All experiments were performed in triplicate.

The higher MIC and MFC values obtained, compared with growth curve assays, likely reflect the effect of the composition of the assay medium, as RPMI contains salts and buffering components that can partially attenuate chitosan activity relative to minimal media conditions.

### 3.3 CFU-based viability assays reveal differential recovery after chitosan exposure

To further evaluate whether growth inhibition correlated with loss of cell viability, CFU-based survival assays were performed across a range of chitosan concentrations. In these experiments, cells were exposed to chitosan in a buffered 10 mM MES-TEA solution at pH 5.0, to avoid potential protective effects of nutrient-rich media.

In *S. cerevisiae*, cell viability was largely preserved across the concentration range tested, with only a moderate reduction in survival observed at 20 and 50 µg/mL chitosan, reinforcing previous observations from growth assays in liquid medium, which indicate that this strain is the most resistant among those analyzed (Figure 3a). In contrast, *C. albicans* displayed a higher susceptibility to chitosan, showing a marked decrease in viability starting at 2 μg/mL, and reaching approximately 1% survival at 20 μg/mL (Figure 3b).

**Figure 3.**
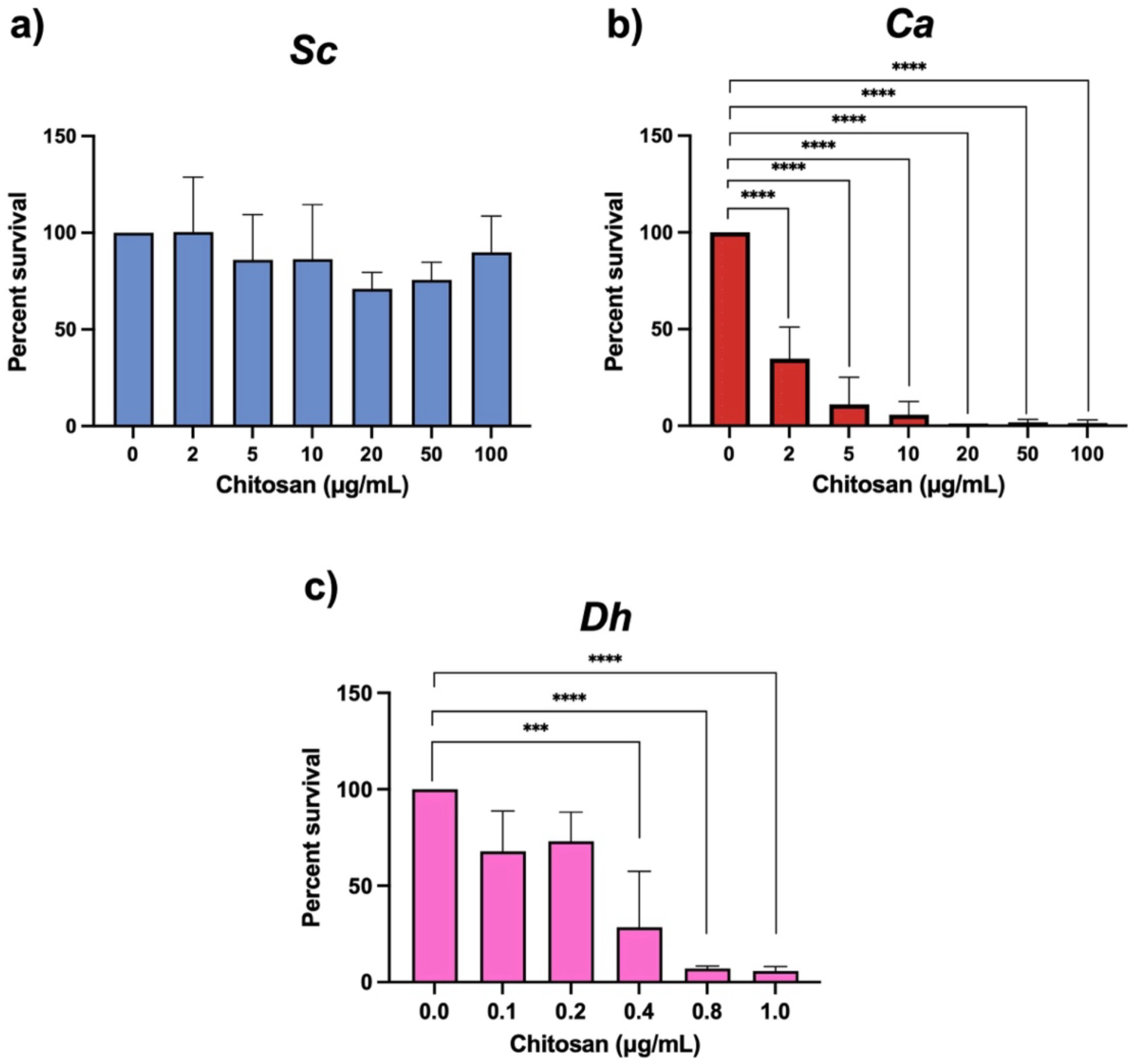
Yeast survival in response to chitosan exposure. Cell survival of *S. cerevisiae* (*Sc*) (a), *C. albicans* (*Ca*) (b), and *D. hansenii* (*Dh*) (c) treated with different concentrations of chitosan was determined by colony-forming units (CFU). Experiments were performed using three biological replicates. Data represent the mean ± SD of three independent biological replicates. Statistical significance was assessed by one-way ANOVA followed by Dunnett’s multiple comparisons test versus the untreated control (0 μg/mL). Differences were considered significant at ***p < 0.001 and ****p < 0.0001.

Notably, *D. hansenii* exhibited the highest sensitivity to chitosan, with a pronounced loss of viability detected at concentrations as low as 0.1 μg/mL, and survival dropping to around 5% at 1 μg/mL (Figure 3c).

### 3.4 Chitosan induces structural alterations in the yeast cell wall and plasma membrane

To examine whether the effects of chitosan observed in the growth and susceptibility assays are related to alterations in cell ultrastructure, transmission electron microscopy (TEM) was used to analyze cells treated with different concentrations, according to preliminary susceptibility estimates.

In untreated cells, all strains, displayed the expected structural organization, showing an intact cell wall and plasma membrane. Particularly, *D. hansenii* exhibited a thicker cell wall compared with the other species. Cells treated with chitosan showed clear structural alterations in cell wall and plasma membrane which differed among the yeast species (Figure 4).

**Figure 4.**
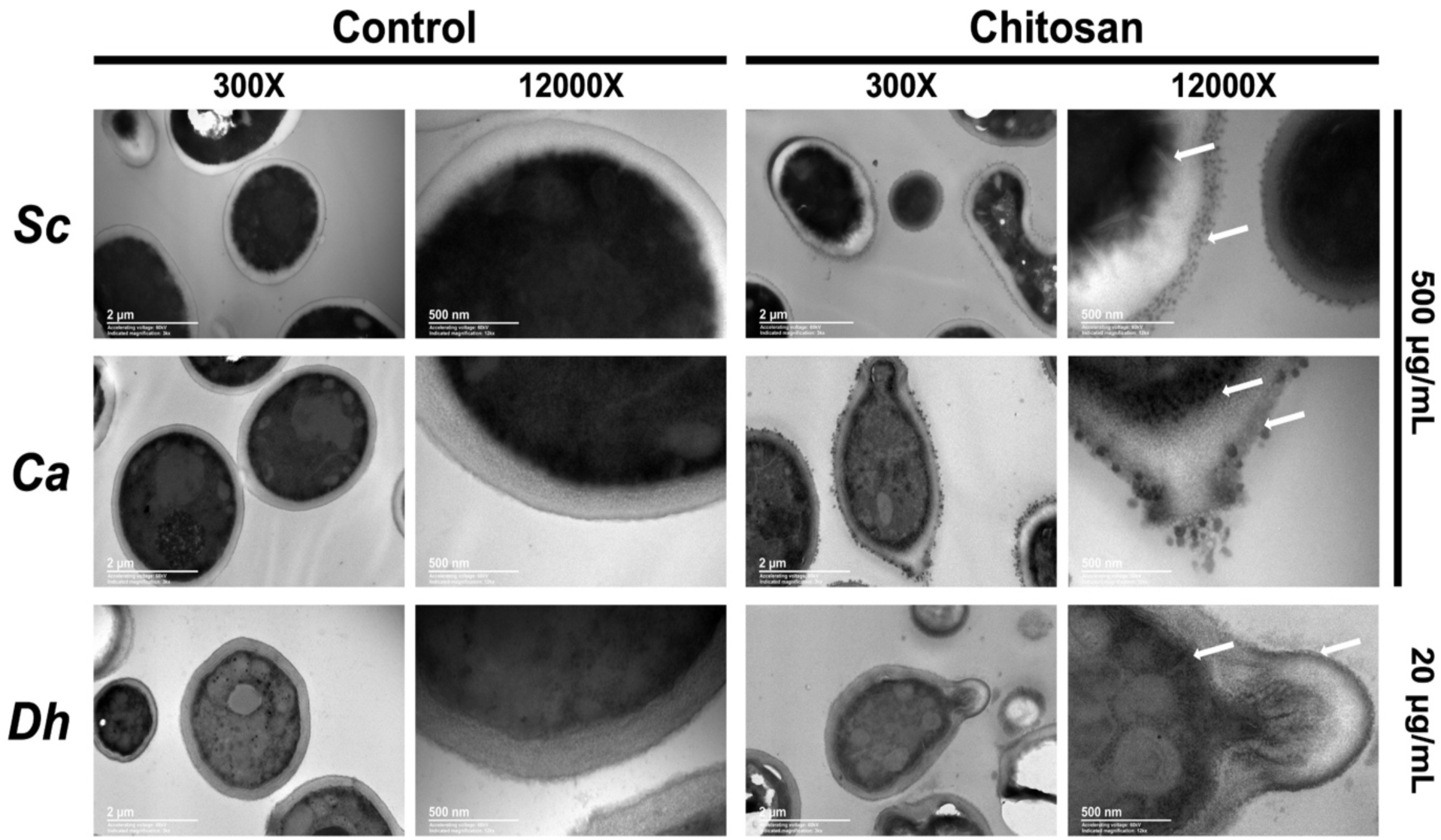
Chitosan-induced alterations of the yeast cell envelope revealed by TEM. Yeast cells treated with chitosan (above the minimum inhibitory concentration [MIC]) were observed by Transmission Electron Microscopy (TEM) and compared with untreated control cells. A concentration of 500 µg/mL was used for *S. cerevisiae* (*Sc*) and *C. albicans* (*Ca*), and 20 µg/mL was used for *D. hansenii* (*Dh*). Representative micrographs show control and chitosan-treated cells at low (300x) and high (12,000x) magnifications. Arrows indicate alterations in the cell envelope (membrane and cell wall) associated with ultrastructural damage. Scale bars: 2 μm (300x) and 500 nm (12,000x).

Exposure to high concentrations of chitosan (500 μg/mL) in *S. cerevisiae*, resulted in evident alterations in the plasma membrane such as invaginations and a brighter appearance. The plasma membrane appeared disorganized and discontinuous, with invaginated portions compared to the untreated cells. Meanwhile, the cell wall also showed disclosed changes as it looked diffuse and damaged. There was also an evident loss of cell shape, which is related to cell wall damage (Figure 4). *C. albicans* presented damage in the cell wall and plasma membrane, with some invaginations in the membrane and in the case of the cell wall it also looked diffuse, and the cells displayed protrusions, evidencing cellular surface damage (Figure 4).

Evidence of the damage was also visible in *D. hansenii*, in the form of invaginations and protrusions caused by the exposure to chitosan (Figure 4).

### 3.5 Chitosan binds to the cellular surface and changes the cellular surface charge

To evaluate how varying levels of chitosan influence the charge on cell surfaces, we examined its interaction with the surface of yeast cells, as it is recognized to modify the surface charge of these cells. This effect was evidenced by a progressive shift in zeta potential values toward more positive charges as chitosan concentration increased, indicating strong electrostatic interactions between the polycationic polymer and negatively charged components of the cell envelope (Figure 5a). This behavior was observed in all species, although the magnitude of the shift differed, suggesting species-dependent surface properties.

**Figure 5.**
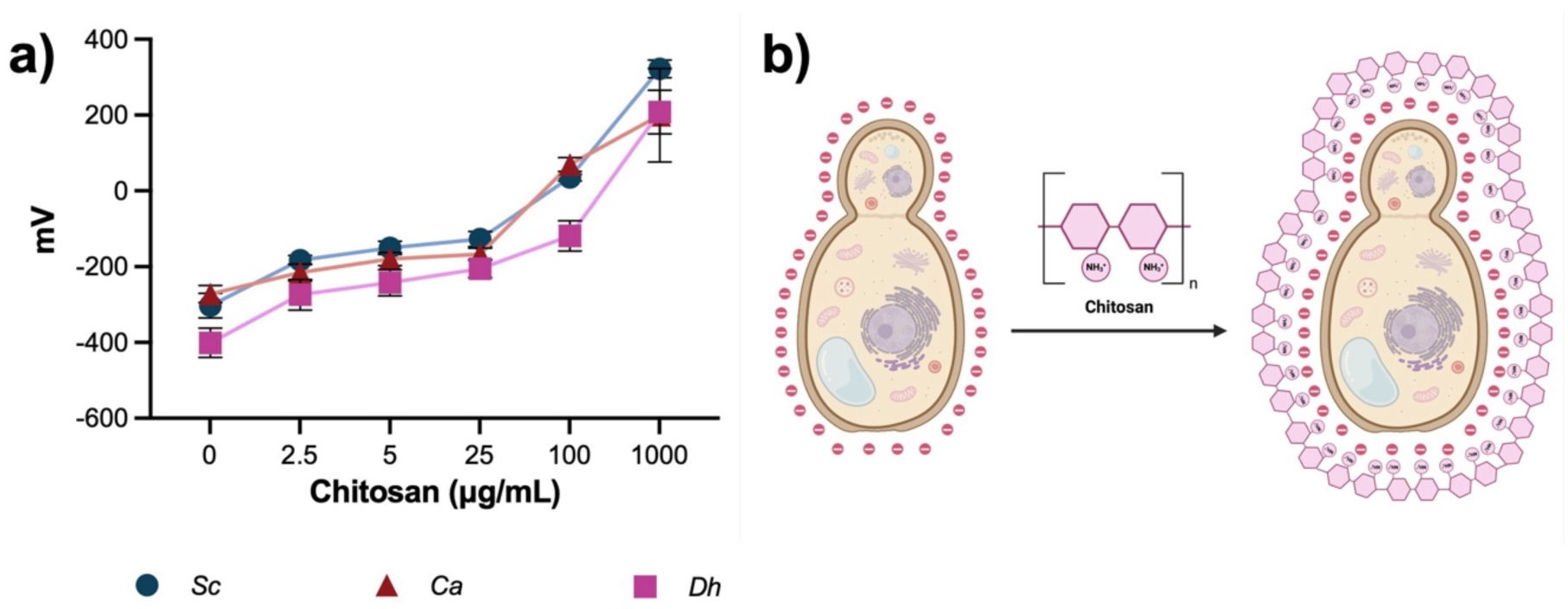
Surface interaction of chitosan with yeast cells. (a) Changes in zeta potential in *S. cerevisiae* (*Sc*), *C. albicans* (*Ca*), and *D. hansenii* (*Dh*) following the sequential addition of increasing concentrations of chitosan, showing a shift from negative to more positive values, consistent with chitosan binding to the cell surface. Data represent mean ± SD (n = 3). Statistical differences between species at each concentration were evaluated by two-way ANOVA followed by Tukey’s multiple comparisons test. (b) Schematic illustration of the electrostatic interaction between protonated chitosan and negatively charged components of the yeast cell envelope, resulting in surface coating.

The proposed mechanism involves electrostatic attraction between protonated amino groups of chitosan and negatively charged residues on the cell surface, resulting in the formation of a polymeric layer surrounding the cells and modifying their physicochemical properties (Figure 5b).

Fluorescence microscopy using FITC–CTS confirmed direct binding of chitosan to the cell surface. In *S. cerevisiae*, untreated cells and cells exposed to FITC alone showed negligible fluorescence, confirming low background signal. Upon exposure to FITC–CTS, fluorescence was detected as early as 10 min, appearing initially as discrete and heterogeneous spots on the cell surface. Over time (1-4 h), fluorescence intensity increased and became more uniformly distributed, indicating progressive accumulation and surface coverage of chitosan. At 24 h, a reduction in fluorescence intensity was observed, possibly associated with cell damage or loss of structural integrity (Figure 6a).

**Figure 6.**
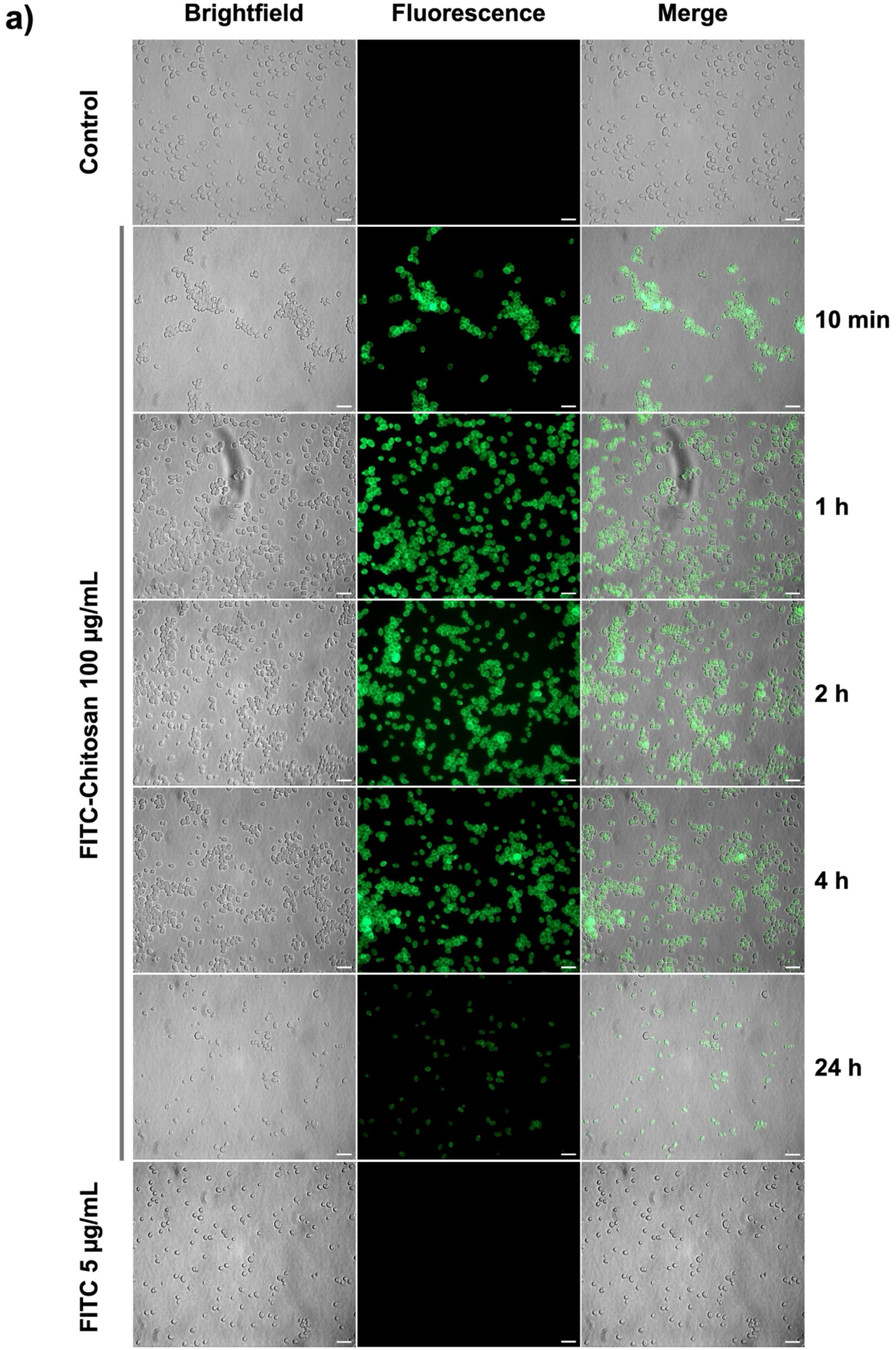
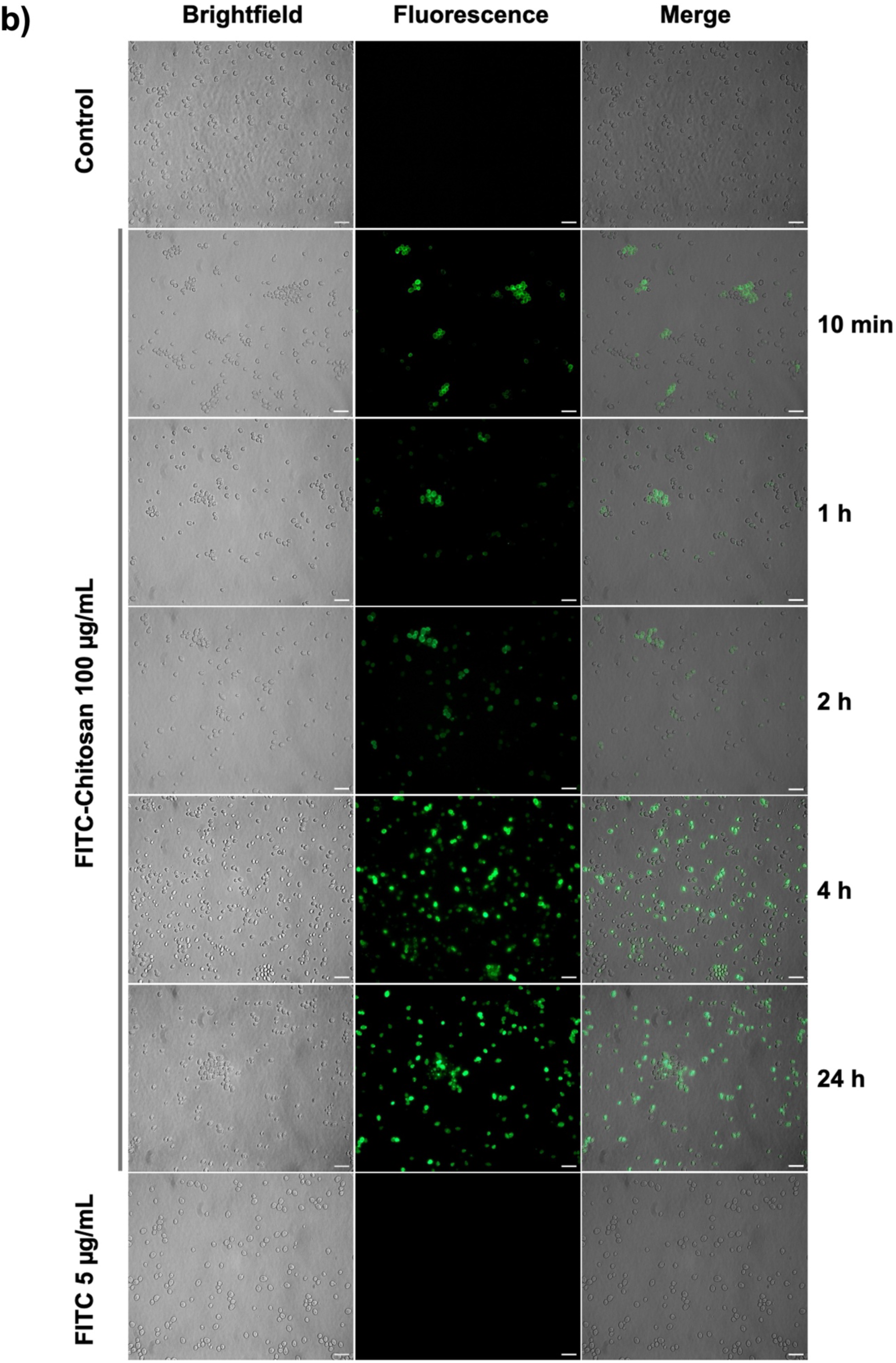
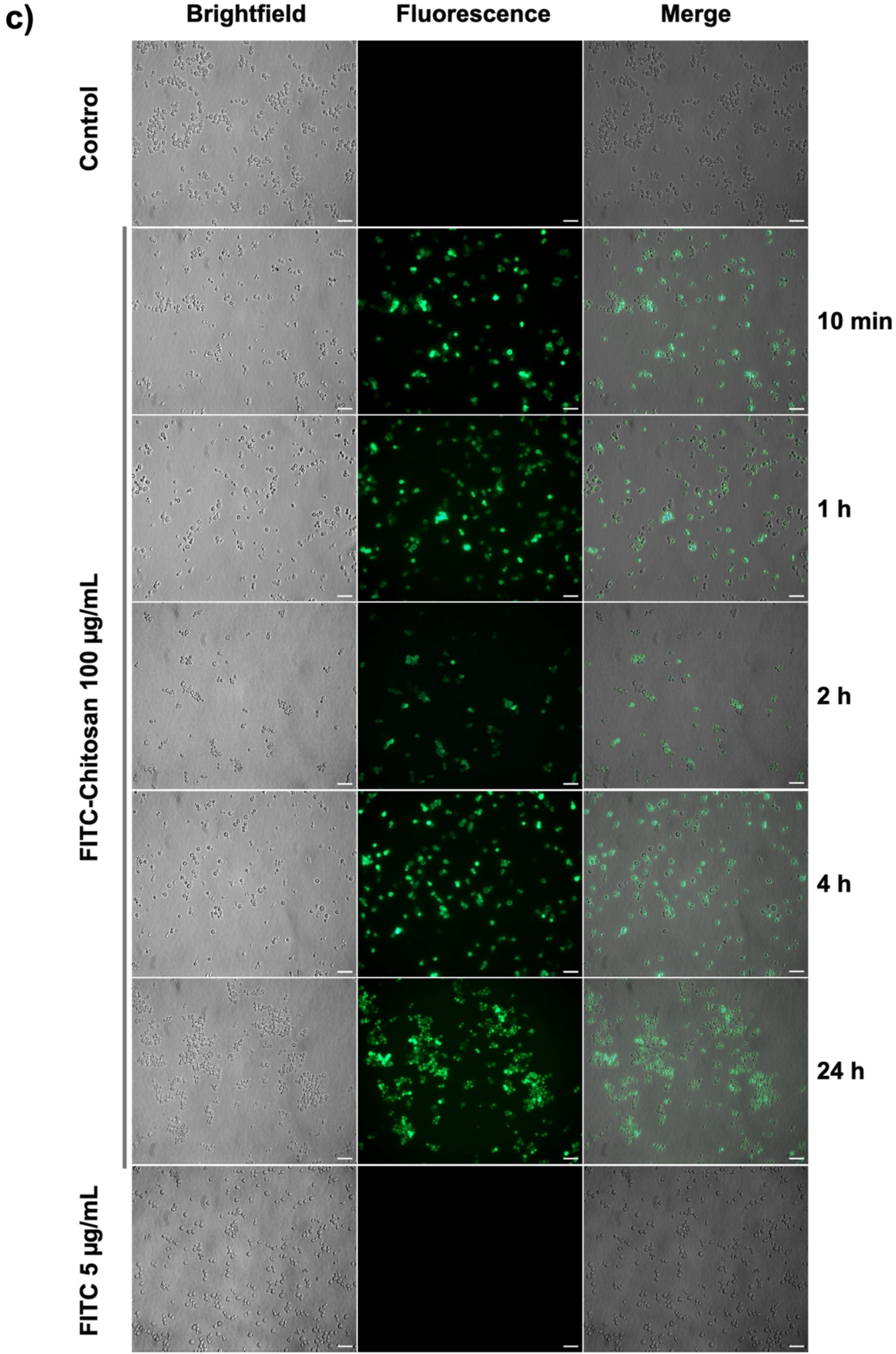
Surface interaction of chitosan with yeast cells. Fluorescence microscopy analysis of *S. cerevisiae (Sc)* (a) *C. albicans (Ca)* (b) and *D. hansenii (Dh)* (c) cells, respectively incubated with FITC–CTS (100 μg/mL) over time (10 min, 1 h, 2 h, 4 h, and 24 h). Brightfield, fluorescence, and merged images are shown. Control cells (untreated) and cells treated with FITC alone (5 μg/mL) exhibited negligible fluorescence. Time-dependent increase in fluorescence intensity and distribution is observed, indicating progressive binding of chitosan to the cell surface. Experiments were performed in three independent biological replicates. Scale bar: 5 μm (100x).

In *C. albicans*, a similar pattern of surface association was observed. Early time points showed localized fluorescence signals, which intensified and spread over the cell surface with increasing incubation time. The fluorescence distribution suggested a strong interaction of chitosan with the cell envelope, although the signal appeared slightly more heterogeneous compared to *S. cerevisiae*, potentially reflecting differences in cell wall composition. At later times, fluorescence persisted but showed signs of redistribution or partial loss (Figure 6b).

In *D. hansenii*, fluorescence signals were detected rapidly, even at early incubation times, indicating a higher affinity or faster interaction with chitosan. The fluorescence intensity increased markedly at 1-4 h, showing a strong and widespread surface signal. Compared to the other species, *D. hansenii* exhibited a more intense and extensive fluorescence pattern, consistent with its higher susceptibility to chitosan. At 24 h, fluorescence was still detectable, although some reduction and heterogeneity were observed, possibly due to cell damage or detachment of the polymer (Figure 6c).

Brightfield images across all species confirmed that during early incubation times, overall cell morphology remained largely intact, while fluorescence images and overlays demonstrated that the signal was predominantly localized at the cell periphery, increasing over time and displaying a heterogeneous distribution among cells, supporting surface binding rather than intracellular accumulation, although some cells exhibited a more diffuse fluorescence pattern at later time points (Figures 6a, 6b, 6c, Supplementary figure 2).

Together, these results demonstrate that chitosan associates strongly with the yeast cell surface in a time-dependent manner, leading to significant alterations in surface charge, likely contributing to downstream effects such as membrane destabilization and permeabilization.

### 3.6 Chitosan alters permeability in a dose-dependent manner in different yeast strains

Observations showed that chitosan affects both the plasma membrane and the cell wall. We therefore evaluated whether treated cells became permeabilized, since permeabilization can disrupt osmotic regulation by altering the balance between the influx and efflux of cellular components and, consequently, compromise cell viability. Several complementary assays were performed to determine whether chitosan induces membrane permeabilization and to assess how this effect depends on its concentration.

First, membrane permeabilization was evaluated using a propidium iodide (PI) assay in a fluorescence plate reader, in cells treated with increasing concentrations of chitosan. Untreated cells were used as a negative control, while heat-killed cells were used as a positive control. In *S. cerevisiae*, more than 50% of cells internalised PI at chitosan concentrations as low as 2.5 μg/mL, and this percentage increazed with higher chitosan concentrations, reaching the highest value at 100 μg/mL. The proportion of permeabilized cells at this concentration appeared higher than that observed in heat-killed cells, which were assumed to be fully damaged and therefore permeable to PI; however, this difference was not statistically significant and should be interpreted as a trend rather than a definitive effect (Figure 7a).

**Figure 7.**
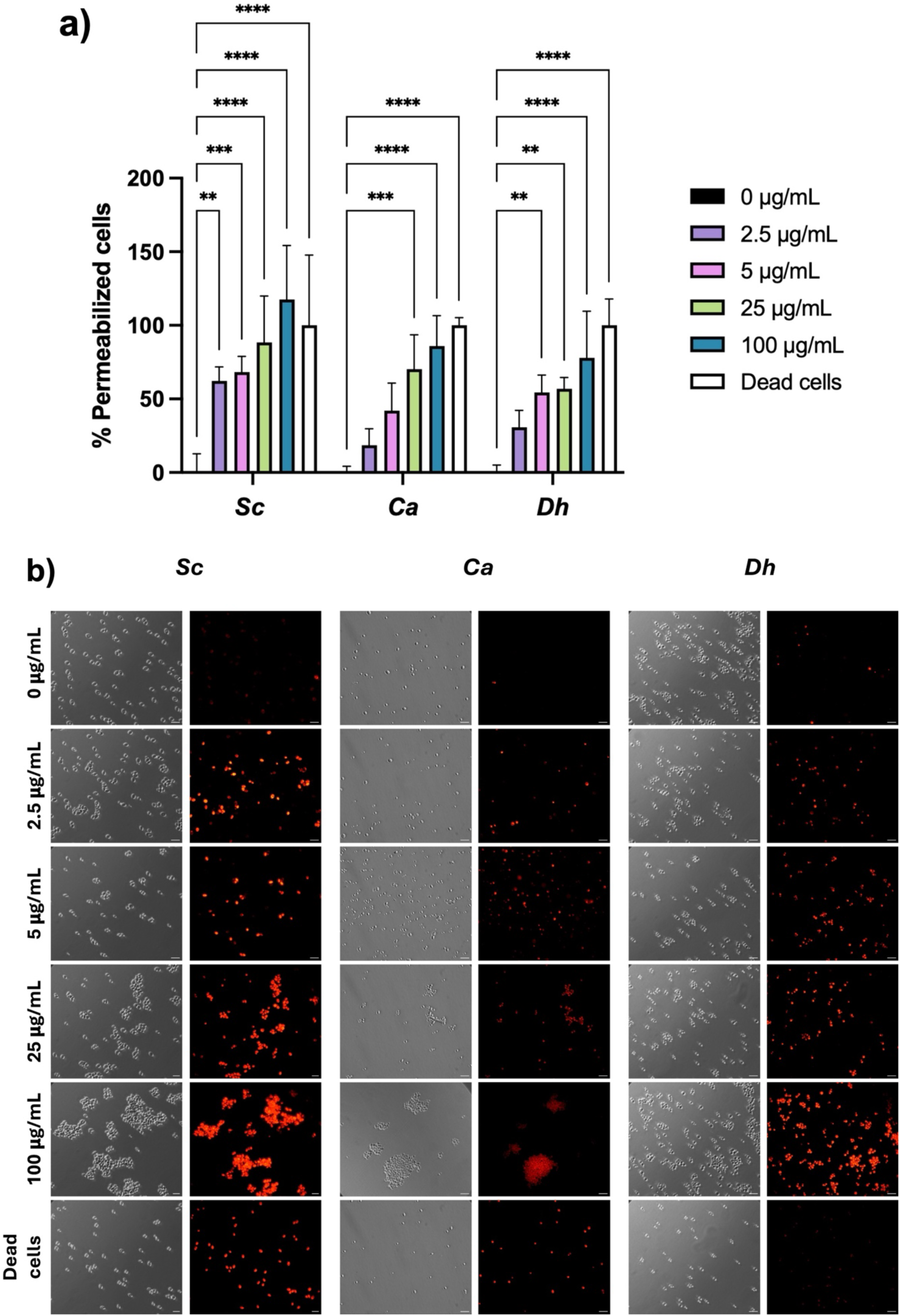
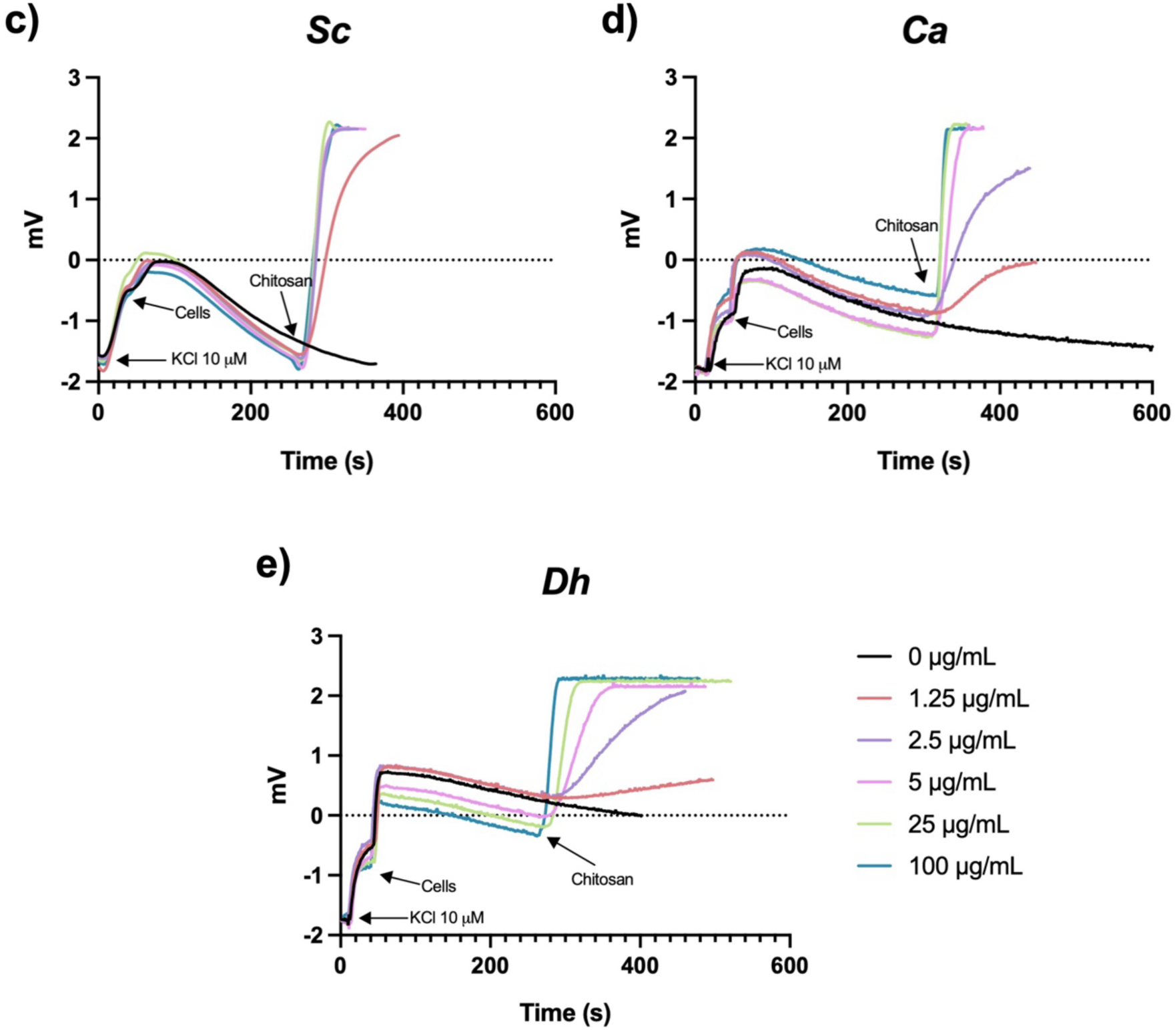
Chitosan Induces Permeabilization in Yeast Cells. (a) Quantification of membrane permeabilization by propidium iodide (PI) uptake in *S. cerevisiae* (*Sc)*, *C. albicans* (*Ca*), and *D. hansenii* (*Dh*) treated with increasing concentrations of chitosan (2.5–100 μg/mL). Data are presented as relative fluorescence intensity. (b) Fluorescence microscopy images of PI uptake under the same conditions. Red fluorescence indicates loss of membrane integrity. Brightfield and fluorescence images are shown for each strain and condition. Untreated cells and heat-killed cells were included as negative and positive controls, respectively. Scale bar: 5 μm. Potassium efflux measurements in *S. cerevisiae* (*Sc*) (c), *C. albicans* (*Ca*) (d), and *D. hansenii* (*Dh*) (e) following exposure to increasing concentrations of chitosan. Changes in extracellular potassium were monitored over time using a selective electrode. A concentration-dependent increase in potassium release was observed, consistent with membrane permeabilization. Representative traces from three independent biological replicates are shown. Data in (a) represent mean ± SD (n = 4). Statistical analysis was performed using two-way ANOVA followed by Tukey’s multiple comparisons test, comparing all treatments to the untreated control within each strain. Statistical significance is indicated as follows: *p < 0.05, **p < 0.01, ***p < 0.001, ****p < 0.0001.

In *C. albicans*, membrane permeabilization was less evident at low chitosan concentrations compared with *S. cerevisiae*. This strain reached more than 50% of permeabilized cells at 25 μg/mL, and even at 100 μg/mL, the degree of permeabilization remained lower than that observed in heat-killed cells. Similar behavior was observed for *D. hansenii*, which exhibited a similar percentage of permeabilized cells as to *C. albicans* across the tested concentrations (Figure 7a). These results were also supported by fluorescence microscopy, where PI staining was detectable at low chitosan concentrations (2.5 μg/mL), with a progressive increase in signal intensity at higher concentrations, in agreement with the relative fluorescence measurements (Figure 7b).

It should be noted that in heat-killed cells, the fluorescence observed by microscopy was not as intense as expected, likely because many cells were disrupted during the boiling process.

To further confirm the permeabilization observed in the PI assay, intracellular potassium efflux was measured in cells treated with the same chitosan concentrations. In this experiment, potassium release was monitored in real time after chitosan addition by means of a selective electrode. In *S. cerevisiae*, a rapid and pronounced potassium efflux was observed at the lowest concentration tested (1.25 μg/mL), indicating a strong membrane permeabilization (Figure 7c); the highest potassium efflux was reached at 2.5 μg/mL. In contrast, *C. albicans* and *D. hansenii* showed slower potassium efflux at the same concentration. *C. albicans* required 5 μg/mL to reach maximal potassium release (Figure 7d), while *D. hansenii* exhibited the slowest potassium efflux among the three strains, although it also reached its highest potassium efflux at 5 μg/mL, this response was slower than that observed in *C. albicans* (Figure 7e).

These results further support the data obtained in the PI assay, showing that membrane permeabilization is not directly related to resistance to chitosan, as the most resistant strain is also the one that became permeabilized at the lowest chitosan concentrations tested in both assays.

### 3.7 *In situ* measurement of enzymatic activity in chitosan permeabilized cells

The previous results highlight the effectiveness of chitosan as a permeabilizing agent across different yeast strains, and we took advantage of this property to measure enzymatic activity *in situ*.

To achieve this, we standardized the appropriate chitosan concentration that would lead to sufficient cell permeabilization while maintaining viability. We performed a test using several concentrations, ranging from the MIC and MFC, to evaluate permeabilization and general dehydrogenase activity of cells, using Alamar Blue as an indicator. We then explored whether chitosan facilitates the entry of small molecules, using NADH as a probe, and observed that permeabilised cells allow NADH to cross the membrane, which does not normally occur in intact cells (Figure 8a). In permeabilised cells, the uptake of NADH is significantly increased compared to the negative control of non-permeabilised cells. We also observed that these permeabilized cells still retained intracellular dehydrogenase enzymes and metabolic activity as compared to a control of dead cells (Figure 8b). This assay allowed us to measure cell permeability, and assess enzymatic activity within the cell. It also enabled us to standardize the amount of chitosan that could be used to permeabilize the cells for an *in situ* enzymatic assay, establishing the ideal concentration for *S. cerevisiae* and *C. albicans* at 125 μg/mL. For *D. hansenii*, the situation was different because it showed very low dehydrogenase activity compared with the other strains, reaching the highest activity at 250 μg/mL (not shown). However, this concentration could not be used, as we previously showed that such high amounts of chitosan affect cellular structure, as observed previously in the TEM analysis. In addition, 10 μg/mL corresponds to the MFC for this strain; therefore, to preserve cell integrity and viability, we determined 5 μg/mL as the optimal concentration. Under these conditions, the structure of the yeast cells was also examined using TEM to confirm their integrity (Supplementary figure 3).

**Figure 8.**
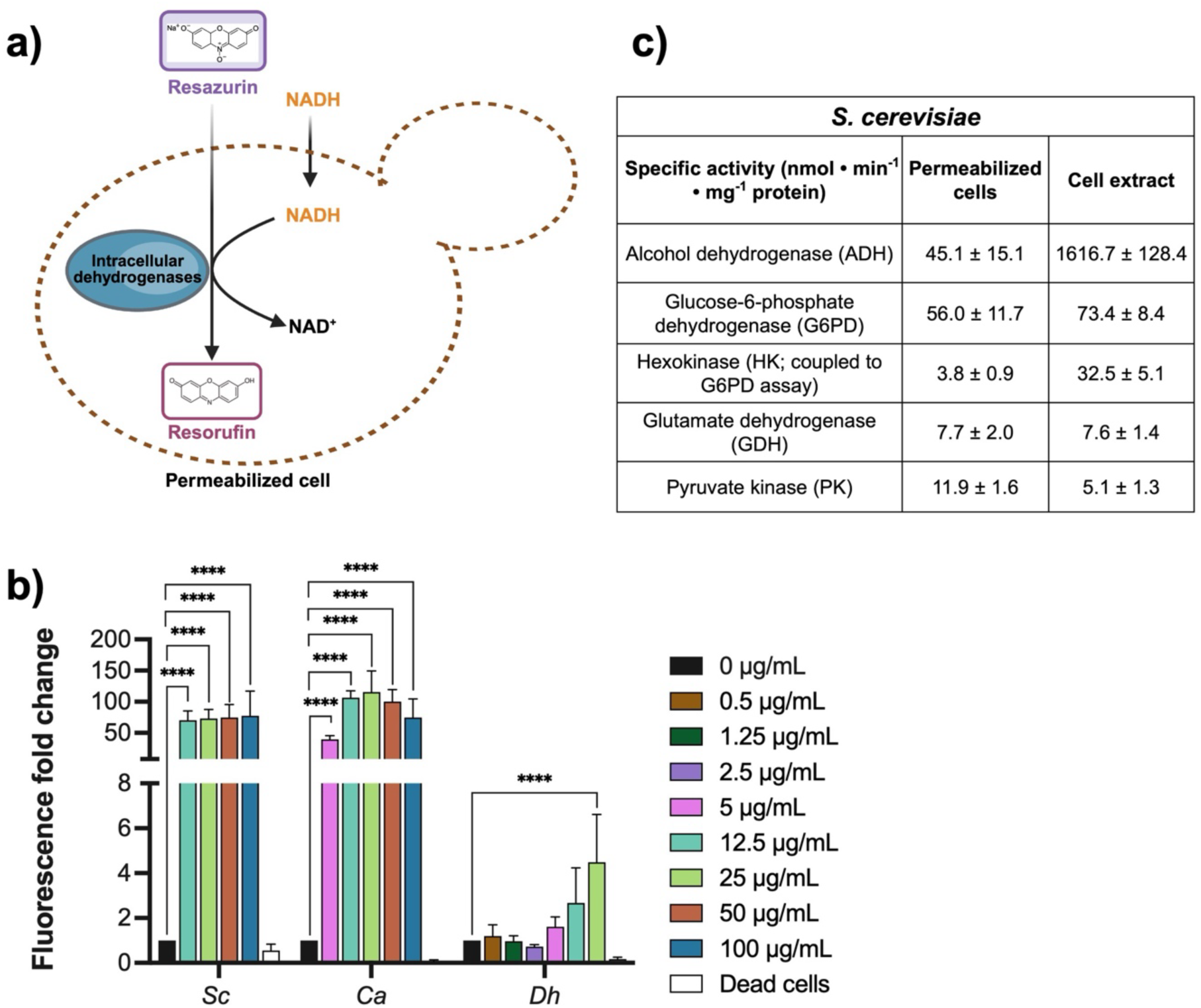
Development of an *in situ* enzymatic assay using chitosan-permeabilised yeast cells. (a) Schematic representation of cell permeabilization and retention of metabolic activity assessed using the Alamar Blue assay. (b) Relative fluorescence changes reflecting intracellular dehydrogenase activity in *S. cerevisiae* (*Sc*), *C. albicans* (*Ca*), and *D. hansenii* (*Dh*) treated with increasing concentrations of chitosan. (c) Specific enzymatic activities measured *in situ* in chitosan-permeabilised *S. cerevisiae* cells and compared with cell-free extracts. Enzyme activities are expressed as nmol min⁻¹ mg⁻¹ protein. Data are presented as mean ± SD from three biological replicates (Alamar Blue: n = 3, with two technical replicates; enzymatic assays: n = 3, with three technical replicates). Statistical analysis was performed using two-way ANOVA followed by Dunnett’s multiple comparisons test versus the untreated control (0 μg/mL) within each strain. Statistical significance is indicated as follows: ****p < 0.0001.

The activity of the enzymes alcohol dehydrogenase (ADH), glucose-6-phosphate dehydrogenase (G6PD), hexokinase (HK), glutamate dehydrogenase (GDH) and pyruvate kinase (PK) was determined only in *S. cerevisiae*, and we successfully detected *in situ* enzymatic activities in chitosan-permeabilized cells and compared them with activities measured in cell-free extracts. Permeabilized cells displayed measurable activities for all the enzymes tested, although at lower levels than those observed in cell extracts.

The enzymatic assays indicated that permeabilised cells displayed an alcohol dehydrogenase (ADH) activity of 45.1 ± 15.1 nmol·min⁻¹·mg⁻¹, in contrast to cell extracts which exhibited a much higher activity of 1,616.7 ± 128.4 nmol·min⁻¹·mg⁻¹ (Figure 8c). In the case of glucose-6-phosphate dehydrogenase (G6PD), permeabilised cells presented an activity of 56.0 ± 11.7 nmol·min⁻¹·mg⁻¹, whereas extracts showed a higher rate of 73.4 ± 8.4 nmol·min⁻¹·mg⁻¹.

The combined activity of G6PD and hexokinase (G6PD/HK) in permeabilised cells was 3.8 ± 0.9 nmol·min⁻¹·mg⁻¹, in comparison to 32.5 ± 5.1 nmol·min⁻¹·mg⁻¹ observed in cell extracts. In permeabilised cells (7.7 ± 2.0 nmol·min⁻¹·mg⁻¹) and cell extracts (7.6 ± 1.4 nmol·min⁻¹·mg⁻¹), the activity of glutamate dehydrogenase (GDH) was found to be almost the same : 7.7 ± 2.0 and 7.6 ±1.4 (nmol·min⁻¹·mg⁻¹) for permeabilised cells and cell extracts respectively.

Furthermore, pyruvate kinase (PK) activity was higher in permeabilised cells (11.9 ± 1.6 nmol·min⁻¹·mg⁻¹) than in cell extracts (5.1 ± 1.3 nmol·min⁻¹·mg⁻¹) (see Figure 8c). Taken together, these results suggest that permeabilization of cells with chitosan enables the evaluation of enzymatic activities within intact cell structures. However, the differences in activity observed between permeabilised cells and cell extracts may reflect variations in enzyme accessibility, stability, or intracellular organisation.

Finally, enzymatic activity was tested in non-permeabilised cells and was found to be negigible for all enzymes assessed, which reinforces the idea that intracellular enzymes are inaccessible without permeabilization (not shown).

## 3 Discussion

The increasing prevalence of fungal infections and resistance to conventional antifungals underscores the practical significance of chitosan as a biocompatible, biodegradable alternative (Alburquenque et al., 2010; Ganan et al., 2019). The present study demonstrates that chitosan exhibits a strain-dependent effect on *S. cerevisiae*, *C. albicans*, and *D. hansenii,* showing that the response to this compound differs among yeast species. Mainly, tolerance to varying chitosan concentrations, as well as the effect of these concentrations on growth rate parameters, indicates that the observed effects are not uniform but rather specific to each strain tested.

Such variability has been previously reported, where the antimicrobial activity of chitosan is influenced by intrinsic cellular, including the composition and organization of the cell wall and plasma membrane (Aranda-Martinez et al., 2016; Lage et al., 2023; Palma-Guerrero et al., 2010).

To start with, the growth patterns identified across the three yeast species studied suggest that chitosan exhibits a fungistatic rather than a fungicidal effect. This is corroborated by the significant decrease in biomass accumulation (AUC), rather than an effect on the viability of cell cultures in *S. cerevisiae* and *C. albicans*, considering also the notable prolongation of the lag phase in *D. hansenii*, while growth remained even at elevated concentrations. Such behavior aligns with a fungistatic mechanism, wherein cellular proliferation is impeded without triggering cell death, as articulated in pharmacodynamic models for antifungal agents like fluconazole, in which growth is reduced but not eliminated even at high concentrations (Venisse et al., 2008).

However, it is crucial to emphasize that growth-based metrics alone (e.g., OD_600_ curves and AUC) are inadequate for definitively differentiating between fungistatic and fungicidal effects. An extensive assessment should encompass supplementary parameters such as viability assays, particularly CFU counts over time, MFC assessment and its correlation with MIC (MFC/MIC ratio), and membrane integrity assays to evaluate loss of viability associated with membrane damage, thereby providing complementary insights into cellular function.

Chitosan tolerance was evaluated though MIC and MCF determination; for this, we based on the CLSI guidelines (Rex, 2008), with some modifications to quantitatively determine the percentage of growth inhibition. The results showed that chitosan exerts a differential strain-dependent effect. In specific *D. hansenii* exhibited the highest sensitivity, with MIC and MCF values several orders of magnitude lower than those in *S. cerevisiae* and *C. albicans*. The high MFC/MIC ratios observed for *S. cerevisiae* and *C. albicans* (>16 and >13, respectively) suggest that, under the tested conditions, chitosan primarily exerts a fungistatic effect. On the other hand, despite *D. hansenii* exhibited the highest sensitivity, its MFC/MIC ratio remained above 4 (12.5), indicating that chitosan still predominantly exhibits a fungistatic rather than a fungicidal effect across all species under the tested conditions. This observation is consistent with the criteria proposed by Hassen (1998), where high MFC/MIC ratios are indicative of growth inhibition without extensive cell killing (Dudiuk et al., 2019; Hassan et al., 1998).

The survival assays are consistent with the strain-dependent effects observed in MIC values and growth in YNBG medium. While *S. cerevisiae* showed only moderate reductions in viability, *C. albicans* and especially *D. hansenii* exhibited a pronounced, concentration-dependent decrease in survival. Notably, *D. hansenii* showed a huge decline even at sub-MIC concentrations, in agreement with its low MIC (0.8 μg/mL), whereas *S. cerevisiae* maintained higher viability, reflecting its greater tolerance.

These findings highlight that, although susceptibility varies markedly among species, the overall mode of action of chitosan is largely growth inhibitory. In line with this, MIC values reported for different fungi and bacteria show considerable variability depending on the physicochemical properties of chitosan, emphasising the importance of its molecular characteristics in determining antimicrobial efficacy (Másson, 2024; Miura et al., 2025). Additionally, differences in cellular architecture contribute significantly to this variability. Fungi, which possess chitin-rich cell walls and distinct membrane lipid compositions, exhibit differential sensitivity to chitosan; for instance, *C. albicans* has been reported to display strain-dependent responses influenced by environmental factors such as pH (Alburquenque et al., 2010; Ganan et al., 2019). Overall, the correlation between MIC, growth inhibition, and survival suggests that chitosan initially acts in a fungistatic manner, with cell death occurring at higher susceptibility levels, particularly in *D. hansenii*.

Previously, chitosan has been demonstrated to exert a significant effect on the cell surface integrity of *C. albicans*, particularly by disrupting the cell wall and cell membrane (Atai et al., 2017). While the exact mechanism of action of chitosan remains a subject of debate, is widely documented that chitosan interacts with the microbial surface, altering cell permeability and interfering with essential transport systems (Atai et al., 2017; Peña et al., 2013). Cell wall integrity also plays a central role in determining sensitivity, as chitosan has been shown to interfere with wall remodeling enzymes and disrupt structural organization, leading to increased vulnerability (Li et al., 2022). In our study, TEM analysis revealed clear alterations in cell envelope structure, including surface irregularities and electron-dense material associated with the cell surface, supporting the idea of direct interaction and structural perturbation of the outer cell layers.

However, our results suggest that structural features, such as wall thickness, are not enough to explain the differences in sensitivity observed among species. Although *D. hansenii* exhibited a thicker cell wall, this did not result in increased resistance. This contrast is significant when compared to *C. albicans ada2Δ* strains, where a reduction in cell wall thickness correlates with increased sensitivity to chitosan and other cell surface stressors (Shih et al., 2019). This interpretation is further supported by studies in *N. crassa* and *Pochonia chlamydosporia*, where variations in chitosan sensitivity resistance are related to cell wall composition and organization (Aranda-Martinez et al., 2016).

Following these observations, one of the first steps needed to explain the effect of chitosan on the yeast species treated, was the determination of its interaction with the cell surface. The observed shift in zeta potential toward more positive values, together with FITC–CTS fluorescence, confirms that electrostatic binding occurs in all three species. This interaction is driven by the attraction between chitosan protonated amino groups and the fungal cell envelope negatively charged components (Lopez-Moya et al., 2019). However, differences in cell wall composition strongly influence the outcome of this interaction. It has been shown that variations in β-glucan organization and wall architecture determine fungal sensitivity to chitosan (Aranda-Martinez et al., 2016). In this context, the higher susceptibility of *D. hansenii* may be related to its cell wall promoting stronger or more persistent chitosan association, thereby facilitating downstream damage. In addition, our fluorescence observations suggested a dynamic interaction process, in which FICT-CTS initially bind to certain regions in the cell surface, followed by progressive distribution across the cell periphery. At longer incubation times, fluorescence was also observed within the cell, suggesting internalization of chitosan. This behavior is consistent with previous reports in *C. albicans* treated with high and low molecular weight chitosan conjugated with FITC (Li et al., 2022).

Importantly, our results indicate that membrane permeabilization does not directly correlate with lethality. *S. cerevisiae* exhibited rapid potassium efflux and early PI uptake at low concentrations but remained the most resistant strain. This suggests that membrane damage, although significant, can be partially tolerated or reversed in this species. Similar observations have been reported in *C. albicans*, where chitosan induces ion leakage without immediate loss of viability, indicating that early permeabilization may represent a sublethal event (Peña et al., 2013). Therefore, the ability of cells to recover from membrane perturbation appears to be a critical determinant of survival.

Notably, *D. hansenii* was the most sensitive species in terms of growth inhibition and loss of viability, despite exhibiting slower potassium efflux and a less pronounced PI uptake compared to *S. cerevisiae*. This apparent discrepancy supports the idea that chitosan toxicity is a multifactorial process involving not only membrane disruption but also alterations in cell wall integrity and cellular physiology (Lopez-Moya et al., 2019; Poznanski et al., 2023; Shih et al., 2019; Suarez-Fernandez et al., 2021). This could be explained by its physiological specialization. *D. hansenii* is a halotolerant yeast with a metabolism highly dependent on ion homeostasis and membrane function (Breuer and Harms, 2006). Small disruptions in membrane integrity or ionic balance may therefore lead to rapid metabolic collapse. Additionally, its response to environmental stress is tightly linked to membrane-associated processes (Sánchez et al., 2008), which could render it more vulnerable to chitosan-induced surface alterations.

Another important factor is that potassium efflux measurements reflect the kinetics of ion release rather than the total extent of cellular damage. Slower efflux in *D. hansenii* may result from differences in intracellular ion pools, membrane transport properties, or structural barriers imposed by the cell wall. Thus, lower potassium release does not necessarily indicate reduced damage. Instead, it is possible that *D. hansenii* undergoes rapid functional impairment before extensive ion leakage becomes detectable. This interpretation is consistent with studies indicating that chitosan can induce oxidative stress, membrane and cell wall remodeling, and metabolic disruption beyond simple permeabilization (Aranda-Martinez et al., 2016; de Azevedo et al., 2024; Krumova et al., 2024; Lopez-Moya et al., 2016; Meng et al., 2020; Shih et al., 2019; Suarez-Fernandez et al., 2021).

At the cellular level, chitosan primarily targets the cell surface, promoting membrane permeabilization and interacting with cell envelope components, which may facilitate its binding and, in some cases, its internalization into the cell.

In contrast, our results revealed distinct permeabilization patterns among the studied strains, suggesting differential susceptibility linked to their physiological and structural characteristics. Moreover, others important factors that should be considered include the physicochemical characteristics of chitosan itself, such as its molecular weight and degree of deacetylation (Poznanski et al., 2023) which are known to influence its interaction with microbial cells.

Taken together, these findings support a model in which chitosan first binds to the cell surface through electrostatic interactions, followed by progressive disruption of the cell wall and plasma membrane. However, the outcome depends on species-specific properties, including cell wall composition, membrane resilience, and metabolic adaptability. The higher sensitivity of *D. hansenii*, despite lower permeabilization indicators, highlights that membrane leakage is not the sole determinant of antifungal activity, but rather one component of a broader and more complex mechanism (Figure 9).

**Figure 9.**
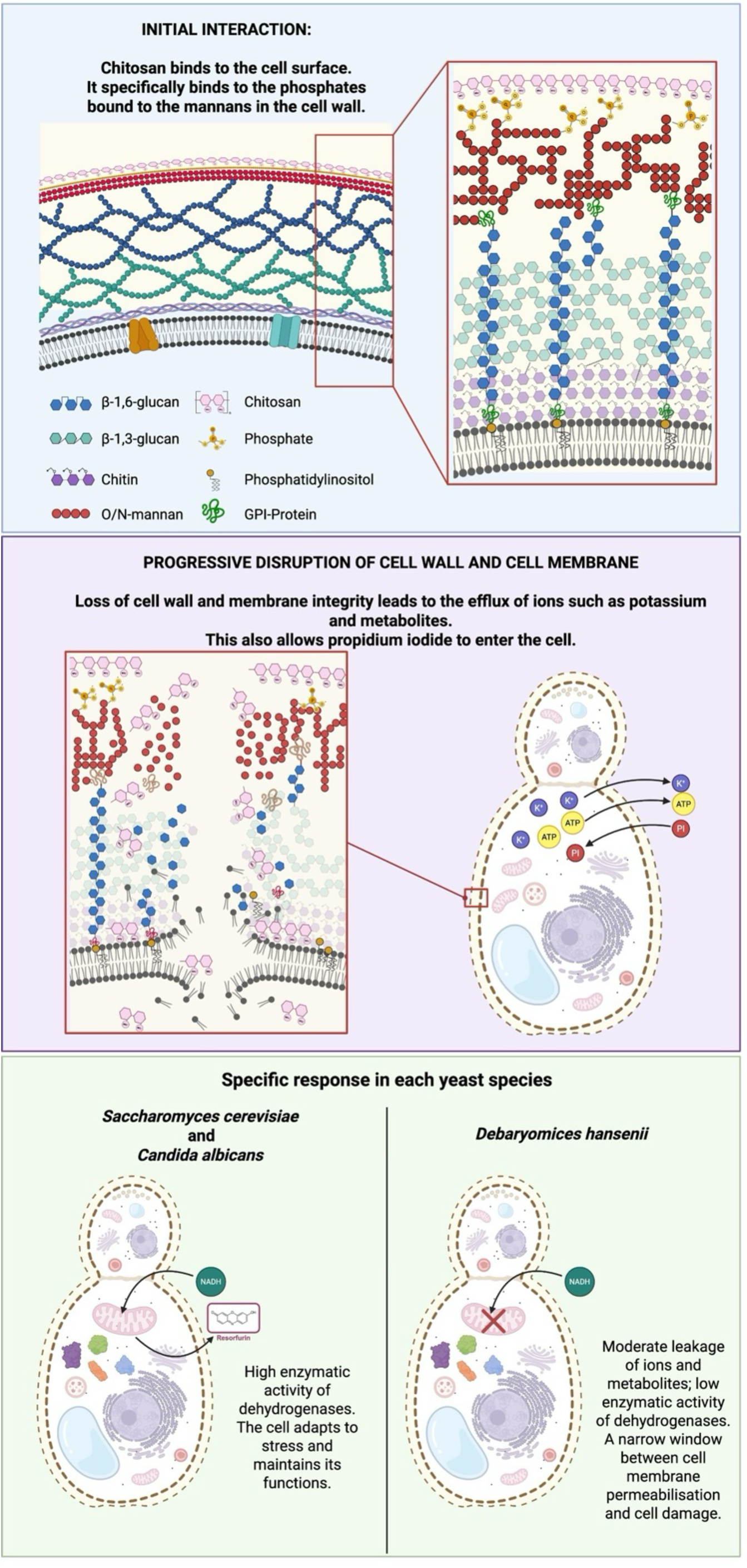
Proposed model of chitosan antifungal activity in yeast. Chitosan binds to the fungal cell surface through electrostatic interactions with mannan-associated negative charges, followed by progressive disruption of the cell wall and plasma membrane, leading to ion leakage and increased permeability. Despite these common effects, the outcome is species dependent. *S. cerevisiae* and *C. albicans* maintain metabolic activity despite membrane damage, whereas *D. hansenii* exhibits reduced metabolic performance despite lower permeabilization indicators. These findings indicate that membrane leakage and metabolic viability are not equivalent, and that chitosan antifungal activity involves a multifactorial, species-specific mechanism.

Finally, the *in situ* enzymatic assays support the idea that permeabilization and metabolic viability are not equivalent. Although permeabilized cells retained measurable enzymatic activity, *D. hansenii* showed reduced metabolic performance compared to the other species. This suggests that chitosan induces a narrower window between reversible permeabilization and irreversible damage in this yeast. In contrast, *S. cerevisiae* appears to better tolerate membrane disruption while maintaining metabolic function, which may explain its higher resistance (Figure 9).

## 5. Conclusion

Chitosan acts as a strain-dependent antifungal agent that primarily targets the yeast cell surface, leading to membrane permeabilization and structural damage. However, susceptibility is not solely determined by the extent of membrane permeabilization, indicating that additional mechanisms such as metabolic disruption and cell wall alterations contribute to its antifungal activity.

The strong correlation between surface binding, changes in zeta potential, and fluorescence patterns confirms that electrostatic interactions play a central role in chitosan activity. Furthermore, the ability of chitosan to induce controlled permeabilization while preserving metabolic function highlights its potential as a useful tool for *in situ* biochemical studies.

Altogether, this work provides new insights into the complex and multifactorial mechanism of action of chitosan and supports its dual application as both an antifungal agent and a methodological tool for studying cellular metabolism in yeast.

## Acknowledgements

The authors thank Dr. Martha Lucinda Contreras-Zentella for technical assistance. We also thank Dr. Natalia Chiquete Félix for support with the fluorescence plate reader, Dr. Roberto Coria Ortega and Dr. Laura Kawasaki Watanabe for assistance with fluorescence microscopy, and Dr. Ruth Rincón Heredia and M.Sc. Rodolfo Paredes Díaz for support with transmission electron microscopy (TEM). The authors are grateful to Mariana Sánchez Maldonado for technical assistance during this project. The authors also thank Sandra Moncada and Javier Gallegos from the Institute’s Library, Juan Barbosa and Ivet Rosas from the Computing Department, and Aurey Galván and Manuel Ortiz from the Maintenance Workshop for their valuable support.

This study was supported by grants IN204321 and IN217924 from the Dirección General de Asuntos del Personal Académico (DGAPA), Universidad Nacional Autónoma de México (UNAM).

The authors wrote and approved the manuscript. Generative AI tools (SciSpace and Jenni AI) were used only for literature search, data extraction, and language editing; all scientific content was generated and critically evaluated by the authors. Mendeley was used for reference management, and Figures 5b, 8a and 9 were created with BioRender.com.

## 6. Author Contributions

Conceptualization: M.A.V., A.P.; Data curation: M.A.V., N.S.S., M.C.; Formal analysis: M.A.V.; Funding acquisition: A.P.; Investigation: M.A.V., N.S.S., M.C., F.P.-G.; Methodology: M.A.V., N.S.S.; Project administration: A.P.; Resources: A.P.; Supervision: A.P.; Validation: M.A.V., N.S.S., M.C., F.P.-G., A.P.; Visualization: M.A.V.; Writing—original draft: M.A.V., A.P; Writing—review & editing: M.A.V., M.C., N.S.S., F.P.-G., A.P. All authors read and approved the final version of the manuscript.

## 7. Funding

This research was supported by grants IN204321 and IN217924 from the Dirección General de Asuntos del Personal Académico (DGAPA), Universidad Nacional Autónoma de México (UNAM).

Minerva Georgina Araiza-Villanueva acknowledges postdoctoral support from DGAPA-UNAM.

## 8. Ethics Statement

All experimental procedures involving *Saccharomyces cerevisiae, Candida albicans* and *Debaryomyces hansenii* were conducted in accordance with the institutional biosafety guidelines of the Instituto de Fisiología Celular, UNAM. The yeast strain used in this study is classified as non-pathogenic and did not require specific ethical approval under current national or international regulations.

## 9. Conflicts of Interest

The authors declare no conflicts of interest.

## Notes

### Competing Interest Statement

The authors have declared no competing interest.

