## Supplementary Tables and Figures for "Effects of Chitosan as a Permeabilizing Agent in Different Yeast Species. Studying Enzymes *in situ*"

**Supplementary Tables 1. AUC of growth in YNBG medium.** Calculated area under the curve (AUC) of growth curves in YNBG medium of *S. cerevisiae*, *C. albicans* and *D. hansenii*. Calculations were performed using GraphPad Prism software with data obtained from growth curves generated from three biological replicates and two technical replicates.

| ***S. cerevisiae*** | | | | | | | | | | | | | | |
| --- | --- | --- | --- | --- | --- | --- | --- | --- | --- | --- | --- | --- | --- | --- |
|  | **5000**  **µg/mL** | **4000**  **µg/mL** | **2500**  **µg/mL** | **2000**  **µg/mL** | **1000**  **µg/mL** | **500**  **µg/mL** | **200**  **µg/mL** | **100**  **µg/mL** | **75**  **µg/mL** | **50**  **µg/mL** | **25**  **µg/mL** | **10**  **µg/mL** | **5 µg/mL** | **0 µg/mL** |
| **Total Area** | 6.483 | 20.68 | 30.77 | 27.17 | 24.87 | 46.17 | 51.24 | 53.94 | 53.81 | 54.51 | 48.43 | 45.55 | 41.07 | 57.81 |
| **Std. Deviation** | 0.4054 | 0.4502 | 0.7682 | 0.7374 | 0.5712 | 0.5272 | 0.4998 | 0.4808 | 0.4313 | 0.4619 | 0.7528 | 1.01 | 1.462 | 0.8163 |
| ***C. albicans*** | | | | | | | | | | | | | | |
|  | **5000**  **µg/mL** | **4000**  **µg/mL** | **2500**  **µg/mL** | **2000**  **µg/mL** | **1000**  **µg/mL** | **500**  **µg/mL** | **200**  **µg/mL** | **100**  **µg/mL** | **75**  **µg/mL** | **50**  **µg/mL** | **25**  **µg/mL** | **10**  **µg/mL** | **5 µg/mL** | **0 µg/mL** |
| **Total Area** | 22.45 | 28.01 | 26.45 | 29.1 | 34.16 | 37.25 | 68.05 | 69.86 | 68.58 | 68.33 | 70.61 | 68.47 | 68.6 | 77.47 |
| **Std. Deviation** | 0.2525 | 0.5536 | 0.9142 | 0.5582 | 2.037 | 1.779 | 0.9768 | 0.9474 | 1.028 | 1.416 | 1.205 | 1.177 | 1.425 | 1.684 |
| ***D. hansenii*** | | | | | | | | | | | | | | |
|  | **100**  **µg/mL** | **50**  **µg/mL** | **20**  **µg/mL** | **10**  **µg/mL** | **5 µg/mL** | **2.5**  **µg/mL** | **1.25**  **µg/mL** | **0.625**  **µg/mL** | **0.375**  **µg/mL** | **0.125**  **µg/mL** | **0 µg/mL** |  |  |  |
| **Total Area** | 3.403 | 1.983 | 2.15 | 1.958 | 4.282 | 16.79 | 30.49 | 37.49 | 35.99 | 41.88 | 41.2 |  |  |  |
| **Std. Deviation** | 0.02812 | 0.07002 | 0.03518 | 0.04458 | 0.3793 | 0.4356 | 0.3343 | 0.3133 | 0.2923 | 0.3103 | 0.2542 |  |  |  |

**Supplementary Table 2. AUC and % Inhibition for MIC determination.** Calculated Area under the curve (AUC) values and corresponding percentage inhibition were calculated from growth curves of *S. cerevisiae*, *C. albicans*, and *D. hansenii* grown in RPMI medium supplemented with increasing concentrations of chitosan. All experiments were performed in triplicate.

| ***S. cerevisiae*** | | | | | | | | | | | | | | | | | | | | | | | | | | | | | | | | | | | | | | | | | | | | |
| --- | --- | --- | --- | --- | --- | --- | --- | --- | --- | --- | --- | --- | --- | --- | --- | --- | --- | --- | --- | --- | --- | --- | --- | --- | --- | --- | --- | --- | --- | --- | --- | --- | --- | --- | --- | --- | --- | --- | --- | --- | --- | --- | --- | --- |
|  | **5000**  **µg/mL** | | **4000**  **µg/mL** | | **2500**  **µg/mL** | | **2000**  **µg/mL** | | **1500**  **µg/mL** | | **1250**  **µg/mL** | | **1000**  **µg/mL** | | **800**  **µg/mL** | | **600**  **µg/mL** | | **400**  **µg/mL** | | **250**  **µg/mL** | | **200**  **µg/mL** | | **125**  **µg/mL** | | **100**  **µg/mL** | | **75**  **µg/mL** | | **62.5**  **µg/mL** | | **50**  **µg/mL** | | **37.5**  **µg/mL** | | **31.25**  **µg/mL** | **25 µg/mL** | **18.75**  **µg/mL** | | **15**  **µg/mL** | | **5**  **µg/mL** | **0**  **µg/mL** |
| **Total Area** | -3.20 | | -1.93 | | -2.54 | | -2.38 | | 0.26 | | 0.56 | | 1.02 | | 1.22 | | 2.22 | | 2.13 | | 1.87 | | 2.52 | | 2.53 | | 2.64 | | 2.39 | | 2.69 | | 3.09 | | 2.28 | | 2.54 | 3.42 | 3.00 | | 3.52 | | 3.60 | 4.70 |
| **Std.**  **Dev** | 3.22 | | 1.62 | | 2.26 | | 2.71 | | 0.81 | | 0.36 | | 1.60 | | 2.11 | | 1.01 | | 0.19 | | 0.30 | | 0.75 | | 0.38 | | 0.73 | | 0.44 | | 0.90 | | 1.14 | | 0.65 | | 0.74 | 1.04 | 0.99 | | 0.86 | | 0.47 | 0.91 |
| **% Inhib.** | 168.22 | | 141.03 | | 154.01 | | 150.73 | | 94.55 | | 88.14 | | 78.18 | | 74.09 | | 52.73 | | 54.62 | | 60.25 | | 46.38 | | 46.12 | | 43.71 | | 49.11 | | 42.73 | | 34.29 | | 51.54 | | 45.83 | 27.20 | 36.04 | | 24.99 | | 23.31 | 0.00 |
| ***C. albicans*** | | | | | | | | | | | | | | | | | | | | | | | | | | | | | | | | | | | | | | | | | | | | |
|  | **5000**  **µg/mL** | | **4000**  **µg/mL** | | **2500**  **µg/mL** | | **2000**  **µg/mL** | | **1500**  **µg/mL** | | **1250**  **µg/mL** | | **1000**  **µg/mL** | | **800**  **µg/mL** | | **600**  **µg/mL** | | **400**  **µg/mL** | | **250**  **µg/mL** | | **200**  **µg/mL** | | **125**  **µg/mL** | | **100**  **µg/mL** | | **75**  **µg/mL** | | **62.5**  **µg/mL** | | **50**  **µg/mL** | | **37.5**  **µg/mL** | | **31.25**  **µg/mL** | **25 µg/mL** | **18.75**  **µg/mL** | | **15**  **µg/mL** | | **5**  **µg/mL** | **0**  **µg/mL** |
| **Total Area** | -1.38 | | 1.62 | | 1.48 | | 5.84 | | 5.75 | | 5.90 | | 4.03 | | 0.41 | | 5.00 | | 4.73 | | 9.42 | | 6.49 | | 9.03 | | 5.93 | | 6.08 | | 4.39 | | 6.73 | | 6.38 | | 8.22 | 9.13 | 8.71 | | 9.47 | | 8.86 | 12.69 |
| **Std.**  **Dev** | 3.35 | | 4.12 | | 9.74 | | 3.75 | | 2.27 | | 2.90 | | 2.49 | | 0.22 | | 2.71 | | 0.75 | | 1.61 | | 4.23 | | 0.18 | | 6.12 | | 6.35 | | 4.24 | | 4.42 | | 5.53 | | 5.62 | 6.23 | 5.91 | | 6.32 | | 5.94 | 3.88 |
| **% Inhib.** | 110.90 | | 87.24 | | 88.30 | | 53.98 | | 54.71 | | 53.50 | | 68.25 | | 96.77 | | 60.58 | | 62.75 | | 25.76 | | 48.85 | | 28.83 | | 53.26 | | 52.09 | | 65.40 | | 46.98 | | 49.72 | | 35.21 | 28.02 | 31.40 | | 25.40 | | 30.17 | 0.00 |
| ***D. hansenii*** | | | | | | | | | | | | | | | | | | | | | | | | | | | | | | | | | | | | | | | | | |  |  |  |
|  | | **100**  **µg/mL** | | **50**  **µg/mL** | | **20**  **µg/mL** | | **10**  **µg/mL** | | **6.67**  **µg/mL** | | **5**  **µg/mL** | | **4**  **µg/mL** | | **3**  **µg/mL** | | **2.5**  **µg/mL** | | **2**  **µg/mL** | | **1.5**  **µg/mL** | | **1.25**  **µg/mL** | | **1**  **µg/mL** | | **0.75**  **µg/mL** | | **0.625**  **µg/mL** | | **0.5**  **µg/mL** | | **0.375**  **µg/mL** | | **0.125**  **µg/mL** | | **0.0694**  **µg/mL** | | **0**  **µg/mL** | |  |  |  |
| **Total Area** | | -0.30 | | 0.61 | | 0.51 | | 0.49 | | 0.54 | | 0.22 | | 0.23 | | 0.09 | | 0.60 | | 0.00 | | 2.20 | | 3.61 | | 4.12 | | 4.92 | | 5.34 | | 5.25 | | 5.39 | | 7.64 | | 7.66 | | 8.29 | |  |  |  |
| **Std.**  **Dev** | | 0.19 | | 0.40 | | 0.48 | | 0.43 | | 0.36 | | 0.32 | | 0.36 | | 0.39 | | 1.03 | | 1.04 | | 1.47 | | 1.06 | | 0.94 | | 1.48 | | 0.91 | | 1.42 | | 1.46 | | 1.88 | | 1.88 | | 1.43 | |  |  |  |
| **% Inhib.** | | 103.57 | | 92.63 | | 93.86 | | 94.09 | | 93.52 | | 97.36 | | 97.25 | | 98.91 | | 92.79 | | 100.05 | | 73.41 | | 56.46 | | 50.33 | | 40.70 | | 35.62 | | 36.65 | | 35.00 | | 7.86 | | 7.65 | | 0.00 | |  |  |  |

**Supplementary Figures**

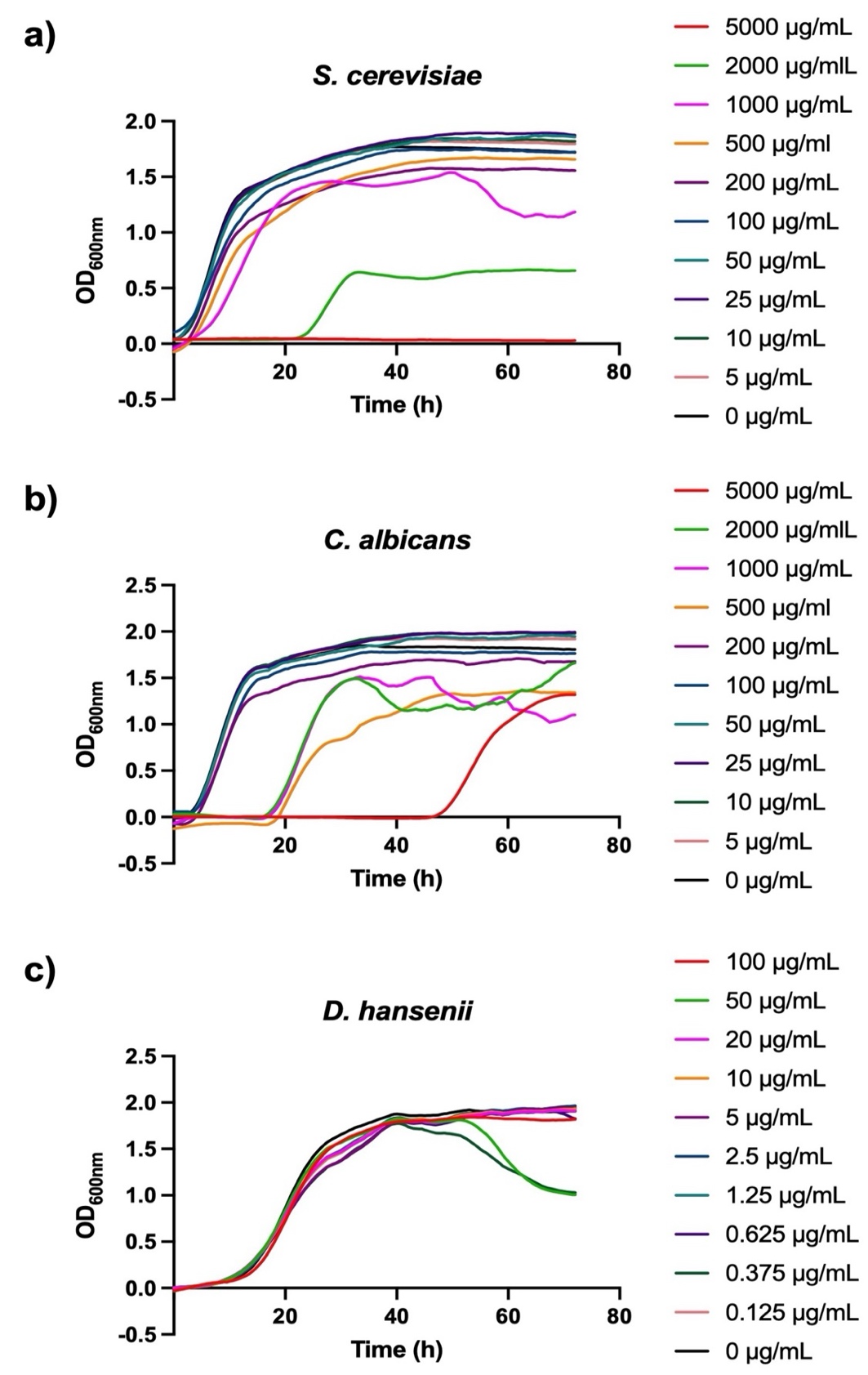

**Supplementary Figure 1. Growth curve in YPD medium.** Growth curves of *S. cerevisiae* (a), *C. albicans* (b), and *D. hansenii* (c) in YPD medium supplemented with various concentrations of chitosan. The curves represent the mean of three biological replicates, each with two technical replicates.

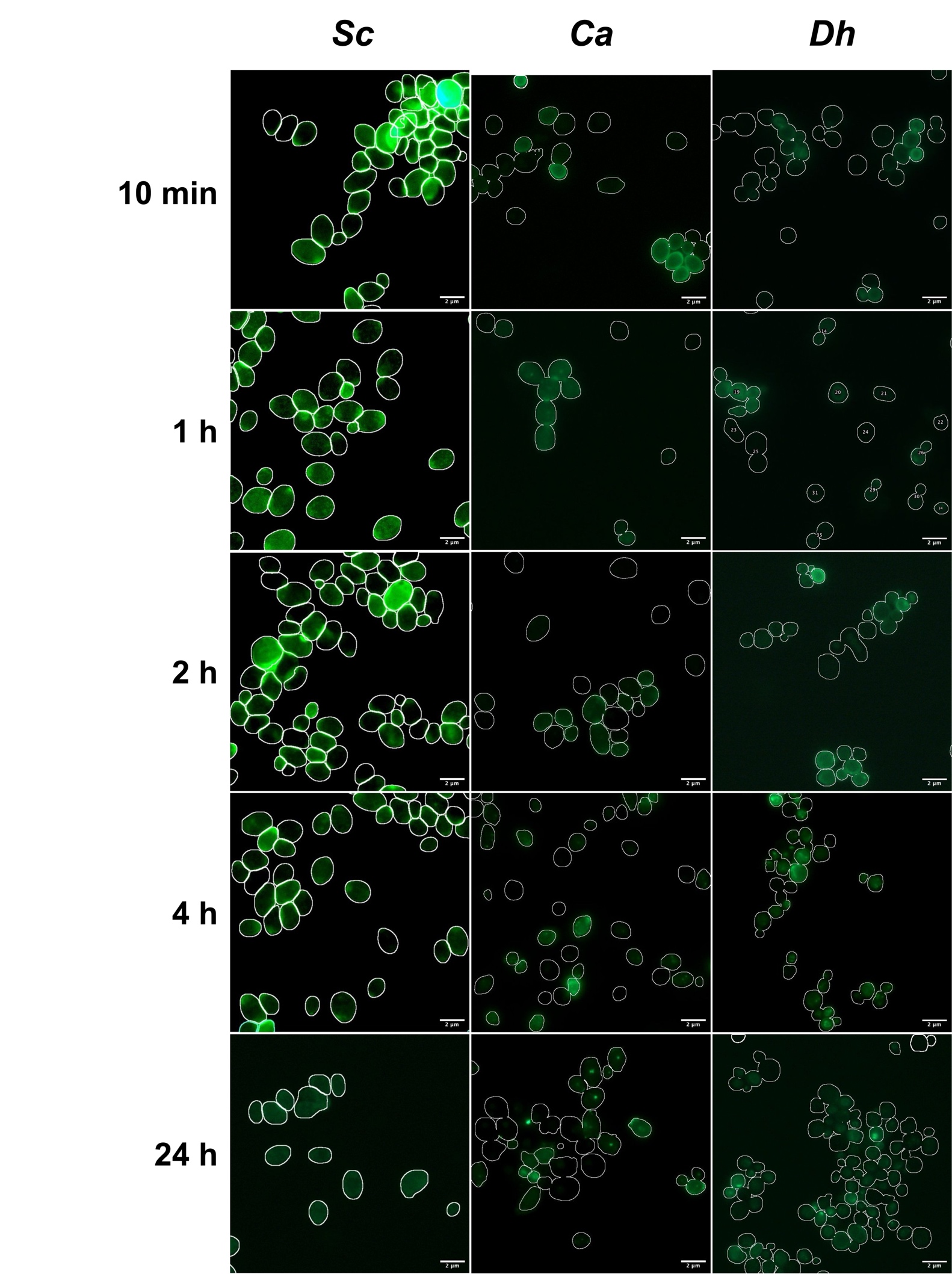

**Supplementary Figure 2. Time-dependent binding of FITC–CTS to yeast cells.** Fluorescence microscopy images showing the temporal dynamics of FITC–CTS association with *S. cerevisiae* (*Sc*) *C. albicans* (*Ca*) and *D. hansenii* (*Dh*) cells at 10 min, 1, 2, 4, and 24 h. Fluorescence outlines the cell periphery, indicating surface binding of chitosan, with intensity increasing at early time points and decreasing at prolonged incubation. Scale bar: 2 μm (100x).

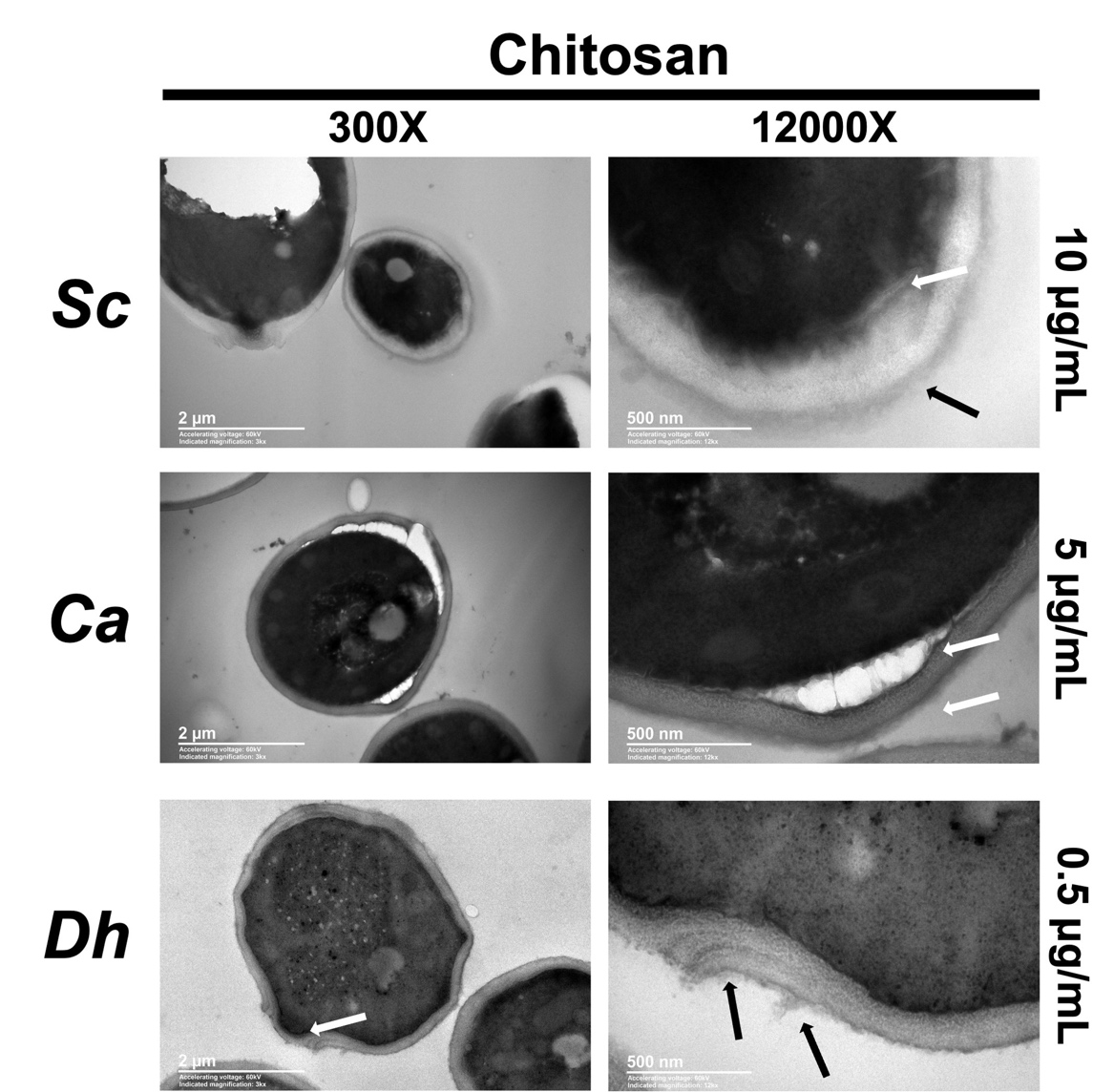
**Supplementary Figure 3. Ultrastructural alterations induced by low concentrations of chitosan in yeast cells**. Transmission electron microscopy (TEM) images of *S. cerevisiae* (*Sc*), *C. albicans* (*Ca*), and *D. hansenii* (*Dh*) exposed to chitosan at the indicated concentrations. Images are shown at 300x (left) and 12,000x (right). Chitosan-treated cells exhibit alterations at the cell surface, including electron-dense material deposition, membrane irregularities, and structural disruptions (arrows). Scale bars: 2 μm (300x) and 500 nm (12,000x).
